# A HigB-like toxin promotes non-replicating *Salmonella* inside macrophages by inhibiting ribonuclease III

**DOI:** 10.1101/2024.07.09.602740

**Authors:** Soomin Choi, Yong-Joon Cho, Seungwoo Baek, Eunna Choi, Yoon Ki Kim, Eun-Jin Lee

## Abstract

Many bacteria are often resistant to antibiotic treatment because they can slow down their growth rate, thereby attenuating the drug’s effectiveness. A similar growth-rate control is observed in pathogens that infect and persist inside their hosts. The bacterial toxin-antitoxin (TA) system serves as a non-heritable phenotypic switch by slowing down growth through the expression of a toxin component. Here, we investigated a HigB-like type II toxin from the intracellular bacterial pathogen *Salmonella* Typhimurium. Unlike other HigB-like toxins that cleave ribosome-bound mRNAs, it does not exhibit endoribonuclease activity. Instead, it inhibits ribonuclease III, which mediates the initial cleavage for rRNA processing, by directly binding to the dsRNA-binding domain of RNase III, thereby decreasing ribosome assembly and bacterial growth. Given that the formation of HigB-like toxin-mediated non-replicating *Salmonella* within macrophages is RNase III-dependent, persister formation by inhibiting RNase III is a newly identified strategy for pathogens to survive within host cells.

## INTRODUCTION

Many bacteria control their growth rate in response to nutritional availability and stress conditions. Often, slow growth is also associated with non-responsiveness to antibiotic treatments due to the antibiotic’s low efficiency in entering the bacterium under these conditions. These phenotypically drug-tolerant bacteria, known as persisters, are induced by various adverse environmental conditions, including nutrient limitation, multiple stress conditions, the presence of antibiotics, and, in some pathogenic bacteria, entry into host cells (Harms et al., 2018). In the intracellular bacterial pathogen *Salmonella* Typhimurium, phagocytosis induces the formation of non-replicating *Salmonella* inside macrophages that are phenotypically similar to antibiotic-mediated drug-tolerant cells (Helaine et al., 2014). Bacterial toxin-antitoxin (TA) systems were identified as one of genetic elements contributing to persister formation. In these systems, the toxin component inhibits major cellular processes, leading to a non-replicating state (Dorr et al., 2010; Harms et al., 2018; Moyed and Bertrand, 1983).

Toxin-antitoxin (TA) systems are widely conserved among bacteria and archaea. They consist of two genetic elements: the toxin component, which inhibits major cellular activities and serves as a non-heritable phenotypic switch for growth rate, and the antitoxin component, which antagonizes the toxin’s activity (Harms et al., 2018).

Depending on the type of antitoxin component and the mode of antagonizing action of antitoxin on the toxin, TA systems are currently categorized into eight different classes (Jurenas et al., 2022). Among these, type II toxin-antitoxin systems have been the most extensively studied over the years. In this system, antitoxins are proteins that directly bind to toxins and neutralize toxin’s various activities. Type II antitoxins also contain DNA-binding domains that bind to the promoter of the type II toxin-antitoxin gene operon and function as repressors, acting as negative regulators even at the transcriptional level (Fraikin et al., 2020). Under certain circumstances, toxins can be activated by preferential degradation of antitoxins or a change in the molecular ratios between toxin and antitoxin, releasing free toxins (Fraikin et al., 2020). Then, these toxins inhibit diverse cellular functions, including cell wall synthesis, DNA replication, cell division, and protein synthesis (Jurenas et al., 2022), many of whose molecular activities and targets are still largely unknown.

Interestingly, type II toxins are predominantly protein synthesis inhibitors (Harms et al., 2018), and thus many type II toxins target protein translation by cleaving mRNAs and tRNAs (RelE, MazF, or VapC) or modifying tRNAs (VapC or TacT) and the elongation factor EF-Tu (Doc) (Harms et al., 2018; Jurenas et al., 2022). Among these, many translation-inhibitory type II toxins encode endoribonucleases that cleave mRNAs in a ribosome-dependent (RelE-type) or ribosome-independent (MazF-type) fashion (Han and Lee, 2020; Harms et al., 2018; Pedersen et al., 2003; Zhang et al., 2003). While MazF cleaves mRNAs by recognizing specific mRNA sequences, RelE binds to the A-site within the ribosome and cleaves ribosome-bound mRNAs in a codon-dependent manner (Pedersen et al., 2003; Zhang et al., 2003). The RelE-type toxins comprise a large RelE superfamily toxin and include RelE, YoeB, YafQ, and HigB (Feng et al., 2013; Maehigashi et al., 2015; Neubauer et al., 2009; Pedersen et al., 2003; Schureck et al., 2016). Although all RelE-superfamily toxins exhibit ribosome-dependent mRNA endoribonuclease activity, they possess different codon cleavage preferences depending on additional interactions between the toxin, ribosomal RNA, and each nucleotide within the ribosomal A-site (Han and Lee, 2020).

*Salmonella* SehA toxin (*<u>S</u>almonella <u>e</u>nterica* <u>H</u>igB-like toxin <u>A</u>) shares homology with HigB toxins from *Escherichia coli, Vibrio cholera,* and *Proteus vulgaris* (Christensen-Dalsgaard and Gerdes, 2006) and belongs to the RelE/HigB superfamily type II toxins (Chimal-Cazares et al., 2020). The *sehA* gene is co-transcribed with the *sehB* gene, which encodes an antitoxin that neutralizes SehA and also functions as a transcriptional repressor of the *sehAB* operon (Chimal-Cazares et al., 2020). Unlike classical toxin-antitoxin systems, the *sehA* toxin gene precedes the *sehB* antitoxin gene (De la Cruz et al., 2013), implying that the ratio of toxin to antitoxin can be higher than that in the classical toxin-antitoxin operon structure. The *sehAB* operon is found only in *Salmonella* strains usually associated with warm-blooded hosts, and a strain with the *sehB* gene deleted exhibited a defect during oral route of *Salmonella* infection (Chimal-Cazares et al., 2020; Lobato-Marquez et al., 2015), suggesting that the function of this operon is required for *Salmonella* pathogenesis.

Based on similarity to HigB from *E. coli* and *P. vulgaris* (40% and 25% identity, respectively), SehA toxin was expected to function as a ribosome-dependent mRNA endoribonuclease, given that both HigB toxins from *E. coli* and *P. vulgaris* cleave ribosome-bound mRNAs at the A-site, thereby inhibiting translation (Christensen-Dalsgaard et al., 2010; Hurley and Woychik, 2009). However, we demonstrate here that despite this similarity, *Salmonella* SehA toxin does not exhibit mRNA endoribonuclease activity. Instead, it binds to ribonuclease III through its dsRNA-binding domain and inhibits RNase III-mediated ribosomal RNA processing, which is the initial step for rRNA processing and ribosome assembly. Thus, SehA-mediated RNase III inhibition results in a decrease in mature rRNA and ribosome assembly, thereby attenuating *Salmonella* growth. We also showed that SehA-mediated growth inhibition *in vitro* and the formation of non-replicating cells within macrophages are RNase III-dependent, revealing a connection between persister formation and rRNA maturation in this pathogenic bacterium.

## RESULTS

### SehA is a bona fide toxin

A previous study reported that the *sehA* (STM14_4845) and *sehB* (STM14_4844) genes encode a toxin and an antitoxin, respectively (Chimal-Cazares et al., 2020) (Figure 1A). To explore the functional role of SehA in *Salmonella* Typhimurium, we cloned the *sehA* gene under the arabinose-inducible promoter. In the presence of arabinose, *Salmonella* expressing SehA significantly slowed down growth compared to vector-expressing cells or non-inducing controls (Figures 1B and 1C). Similarly, a strain lacking the SehB antitoxin showed delayed growth in N-minimal media regardless of Mg^2+^ concentration, presumably due to the accumulation of free SehA proteins (Figures 1D and 1E).

**Figure 1.**
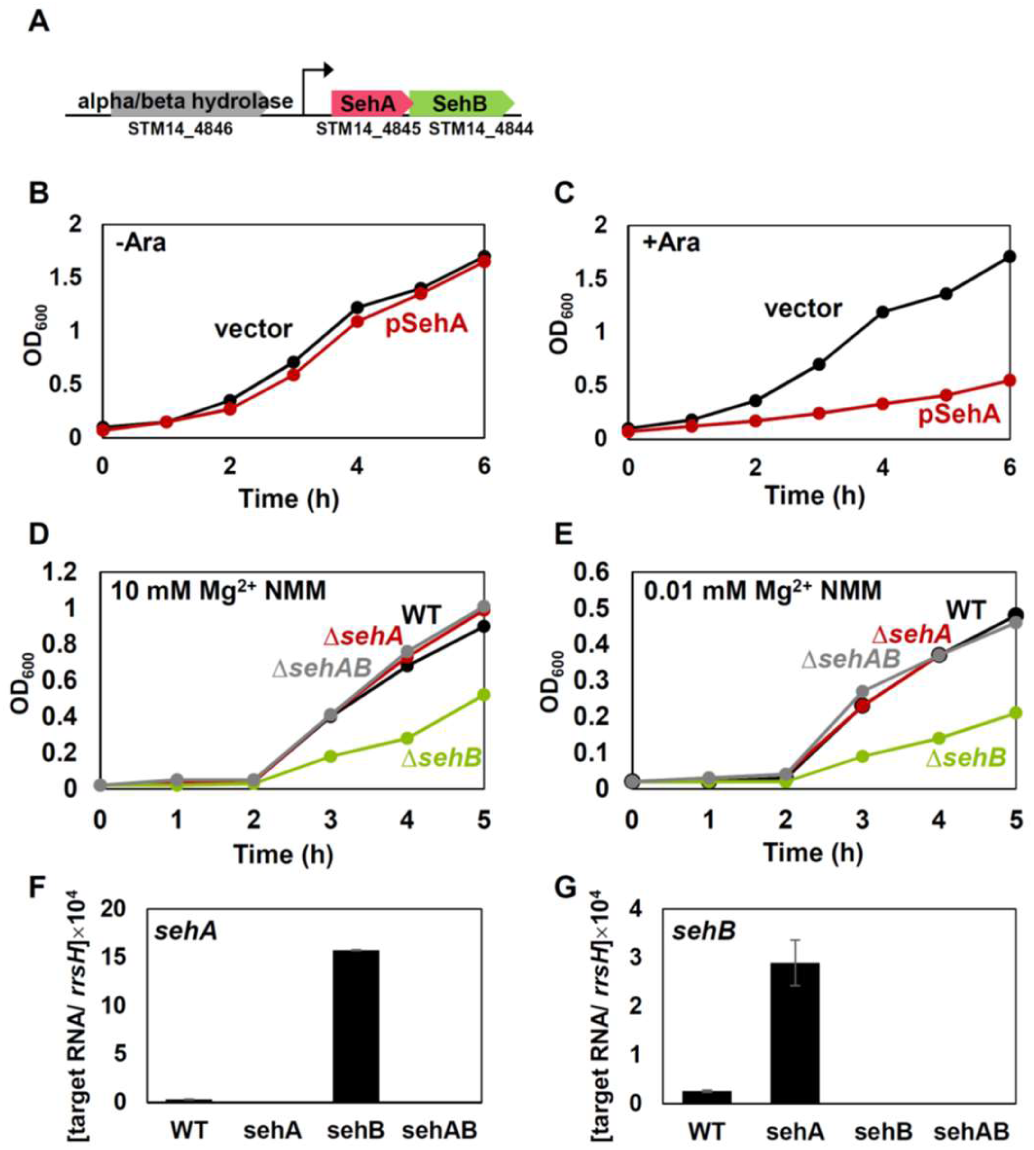
SehA toxin inhibits Salmonella growth. (A) Genetic organization of sehA-sehB toxin-antitoxin operon. (B-C) Growth curves of Salmonella strains expressing SehA (pSehA) or the empty vector (vector) in the presence (C, +Ara) or absence (B, -Ara) of 10 mM arabinose. Bacteria were grown in N-minimal medium containing 0.01 mM Mg^2+^ at 37°C for 6 h with shaking and measured absorbance at OD600 every hour. (D-E) Growth curves of wild-type, the sehA deletion mutant (ΔsehA), sehB deletion mutant (ΔsehB), and a strain deleted both the sehA and sehB genes (ΔsehAB) grown in N-minimal medium containing 10 mM (D) or 0.01 mM Mg^2+^ (E) at 37°C for 5 h with shaking and measured absorbance at OD600 every hour. (F-G) Relative mRNA levels of the sehA (F) and sehB (G) genes in Salmonella strains grown for 3 h in N-minimal medium containing 10 mM Mg^2+^.

Additionally, SehA expression from its native promoter in the absence of SehB appears sufficient to inhibit growth in this medium (Figures 1F and 1G). As controls, strains lacking SehA alone or both SehA and SehB proteins did not exhibit growth defects in the same media (Figures 1D and 1E). These data indicate that SehA is a bona fide toxin.

### Critical residues in SehA toxin are mostly located at the C-terminal α4-helix

When we analyzed the secondary structure of the SehA toxin, we found that the SehA protein consists of three helices (α1-3) followed by three β strands (β1-3) and an additional helix (α4) (Figure 2A). Homology-based protein modeling revealed that α1-3 and α4 helices in SehA are located on the opposite side of the β sheet formed by β1-3 strands, resembling an RNase H-like fold (Figure 2B). This structure modeling also showed that SehA shares homology to HigB in *E. coli* (Yang et al., 2016).

**Figure 2.**
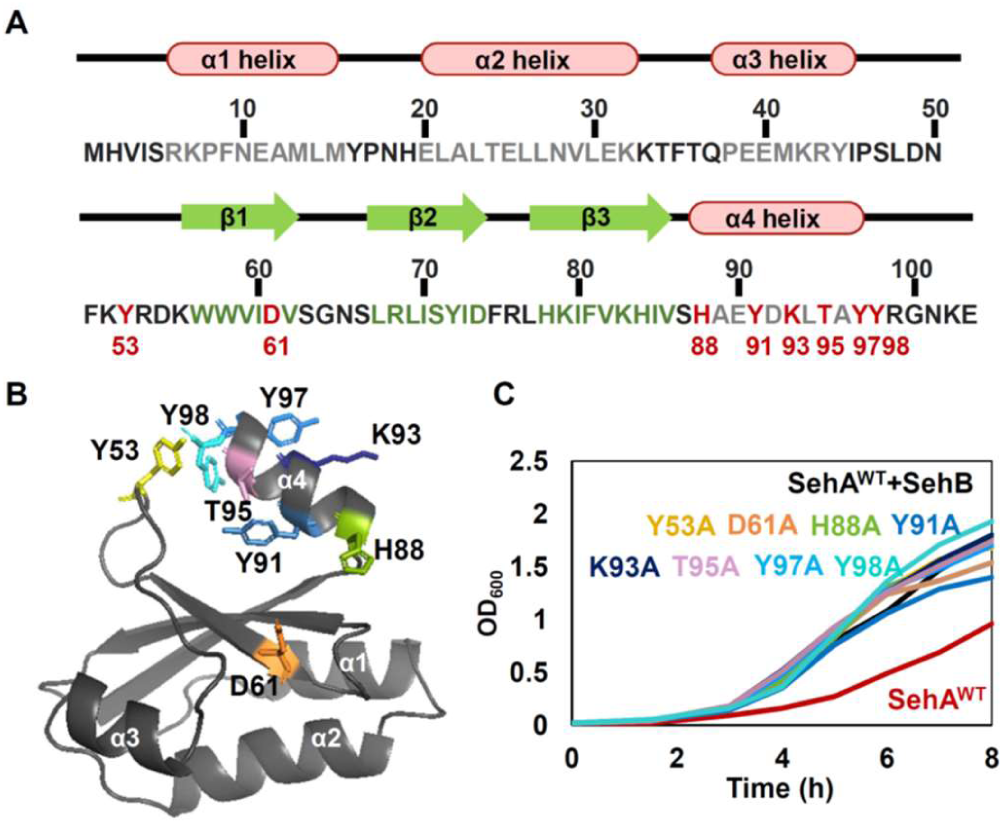
Residues required for SehA toxin activity are located at the additional α4 helix at its C-terminus. (A) Amino acid sequence of the SehA protein from Salmonella enterica serovar Typhimurium 14028s. Schematic illustration of SehA secondary structure prediction is based on the homology modeling using the crystal structure of E. coli HigB (PDB: 5IFG). Four helices and three strands are also indicated as a diagram above the sequence. Sequences corresponding to α-helices and β-strands are colored grey and green respectively. Substituted residues in (C) are indicated and colored red. (B) Homology-modeled structure of the SehA toxin using the crystal structure of E. coli HigB (PDB: 5IFG). Residues required for SehA toxin are mostly located at helix α4. (C) Growth curves of wild-type, the sehB deletion mutant (sehBΔ(155-429)) with the wild-type sehA gene or sehA derivatives with nucleotide substitutions (Y53A, D61A, H88A, Y91A, K93A, T95A, Y97A, or Y98A) grown in N-minimal medium containing 10 mM Mg^2+^ at 37°C for 8 h with shaking and measured absorbance at OD600 every hour.

To determine the key residues required for SehA toxin’s activity, we created chromosomal *sehA* substitution mutants in the *sehB* deletion background to eliminate SehB’s neutralizing effect. Similar to what we observed in Figure 1D, the *sehB* deletion strain with the wild-type *sehA* gene (SehA^WT^) exhibited significantly slower growth compared to the isogenic strain retaining the *sehB* gene (SehA^WT^+SehB) (Figure 2C). SehA variants with Y53A, D61A, H88A, Y91A, K93A, T95A, Y97A, and Y98A substitutions recovered growth in the *sehB* deletion background, indicating that these residues are required for SehA’s toxin activity (Figure 2C). Interestingly, except for the Y53 and D61 residues, all key residues are located at the C-terminal α4 helix (Figure 2B). As controls, two other SehA variants with R54A or K56A substitutions still exhibited growth defects similar to the wild-type SehA, showing that the R54 and K56 residues are dispensable for SehA’s toxin activity (Figure S1).

### SehA toxin alone does not exhibit ribonuclease activity

In the above section, homology-modeled SehA toxin structure revealed that SehA is homologous to HigB proteins from *E. coli* (5IFG; 40% identity) and from *P. vulgaris* (4MCT; 25% identity), which belong to the RelE superfamily toxin that cleaves ribosome-bound mRNAs (Schureck et al., 2016). A previous study also reported that HigB from *P. vulgaris* cleaves ribosome-bound mRNAs at A-rich codons (Hurley and Woychik, 2009). Interestingly, when compared to HigB from *P. vulgaris*, *Salmonella* SehA has an extra C-terminal helix (α4) (Figure S2), exhibiting a three-layer protein fold with α-helices/β-sheet/α-helix (Figure 2B; α1α2α3-β1β2β3-α4). Additionally, critical residues in SehA are clustered in the additional α4 helix (Figure 2), suggesting that SehA toxin might function differently. To explore this, purified GFP-tagged SehA protein was mixed with total RNA extracts to test whether SehA alone has ribonuclease activity *in vitro* (Figure S3). RNA extracts mixed with purified SehA protein did not change rRNA profiles over time (Figure S3), suggesting that purified SehA protein does not exhibit ribonuclease activity *in vitro*. As a control, adding purified SehB proteins to the SehA-containing reaction did not change rRNA profiles either (Figure S3).

Next, to explore differentially expressed genes when SehA toxin is highly expressed by deleting the SehB antitoxin, we analyzed RNA profiles of the wild-type and the *sehB* deletion mutant strains by RNA sequencing. The *sehB* deletion increased *sehA* mRNA levels more than 32-fold (Figure S4), indicating that the *sehB* chromosomal deletion has a SehA-overexpressing effect. Interestingly, SehA overexpression increased mRNA levels of ribosomal protein genes (Figure S4), suggesting that SehA-mediated growth defect might alter ribosome composition. SehA overexpression also downregulated mRNA levels of several genes, most of which are unlikely to be cellular targets for the SehA toxin in inhibiting bacterial growth (Figure S4).

### SehA toxin induces RNA cleavages in the leader regions of rRNA operons

As SehA belongs to the RelE superfamily protein that is predicted as a ribosome-associated endoribonuclease, SehA-mediated growth inhibition could be due to the cleavage of translating cellular target mRNA. Given that RelE-type toxin-mediated mRNA cleavage generates RNA fragments with either 2′-3′ cyclic phosphate or 3′ phosphate at the 3′ end and an RNA fragment with 5′ hydroxylated (OH-) or 5′ monophosphorylated (monoP) group (Feng et al., 2013; Maehigashi et al., 2015; Neubauer et al., 2009) (Figure S5A), we performed a modified RNA sequencing to identify cellular targets of SehA.

To include SehA-generated RNA fragments potentially with a 5′ OH end, the RNA fragments were phosphorylated and ligated to adaptors, followed by sequencing (Figure S5). When we mapped the first nucleotide ligated to the adaptor, the newly appeared reads upon SehA induction, compared to non-inducing control, would correspond to SehA-induced RNA cleavage sites (Figure S5B). Because this method not only detects RNA fragments with a 5′ OH but also with a 5′ monophosphate (5′ OH or monoP RNA), we compared RNA sequencing profiles after directly ligating to the same adaptor, detecting only 5′-monophosphorylated RNA fragments (5′ monoP RNA).

A genome-wide analysis of 5′ OH or monoP RNA ends upon SehA induction revealed that SehA-dependent cleavage sites were not correlated with specific codon or mRNA sequences, and most cleavage sites were enriched in non-coding RNAs (Tables S1 and Table S2). This is surprising given that SehA belongs to the RelE superfamily, which is usually associated with ribosomes and cleaves mRNAs in a codon-dependent manner (Pedersen et al., 2003). Among the non-coding SehA cleavage sites, the most recognizable cleavage sites are located within rRNA operons (Figure 3A). One is located at the -115 nt position relative to the +1 nt of the mature 16 rRNA sequence, and others are located 22 nt after the mature16S rRNA (+1576 in the *rrsH* gene), and 7 nt before the 5′ end and 8 nt after the 3′ end of the mature 23S rRNA (+2049 and +5155 in the *rrlH* gene) sequences (Figure 3B). Among these, SehA-dependent cleavage effects at read positions corresponding to -115 and +2049 appear to be prominent, and these cleavage patterns were almost identical when we compared to 5′-monophosphorylated RNA libraries generated by directly ligating adaptor without phosphorylation (Figure 3C). This suggests that, contrary to our expectations, SehA-mediated RNA cleavage generates RNA fragments with a 5′ monophosphate.

**Figure 3.**
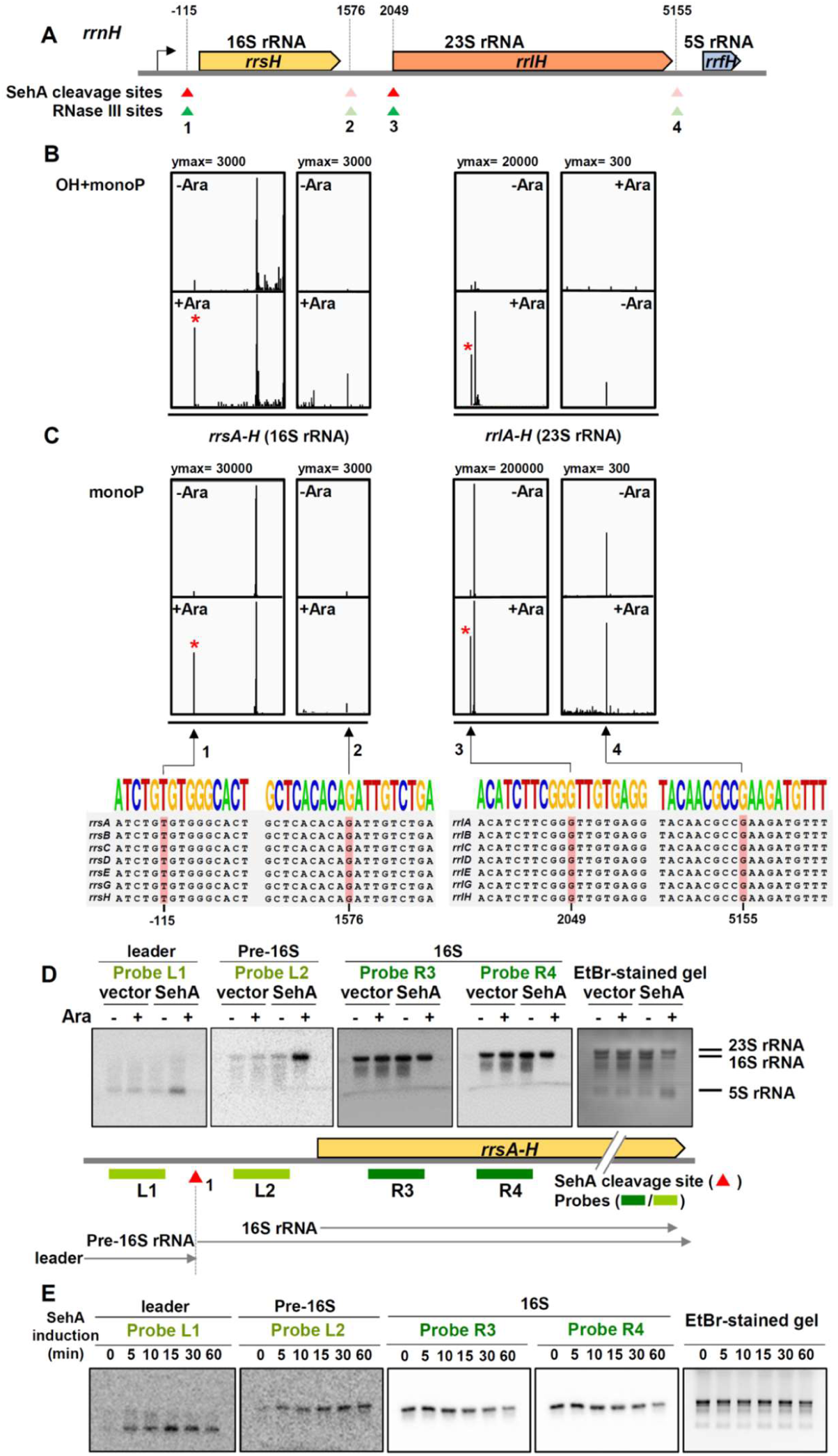
SehA toxin induces RNA cleavages in rRNA operons. (A) Schematic view of Salmonella rrnH 16S-23S-5S rRNA operon. Mature rRNA sequences are represented as colored boxes. The nucleotide positions are listed above relative to the 5’ end of the mature 16S rRNA. SehA toxin cleavage sites are shown in red triangles and RNase III cleavage sites are shown in green triangles. (B-C) SehA-mediated RNA cleavage profiles of 16S rRNA and 23S rRNA genes. (B) 5’ OH- and monoP-RNA profiles. (C) 5’ monoP-RNA profiles. SehA-dependent reads are marked with asterisk (*). The SehA cleavage sites and neighboring sequences are all conserved in seven 16S rRNA and 23S rRNA genes. The nucleotide positions correspond to the rrsH and rrlH rRNA genes in (A). The nucleotide positions in other rRNA operons are listed in Figure S6. Note that the vertical axis is adjusted to visualize SehA-dependent cleavage sites in peaks 1 and 3. (D) Northern blot analysis of rRNAs extracted from wild-type Salmonella strains expressing SehA toxin (SehA) or the empty vector (vector) in the presence (+) or absence (-) of 10 mM L-arabinose. Total RNAs isolated from strains listed above were separated on a MOPS-agarose gel and stained with ethidium bromide (EtBr, far right). The gel was transferred on a membrane and then the membrane was hybridized with probes L1, L2, R3, and R4. The mature 16S rRNA and the precursors (leader and pre-16S) are indicated below. (E) Time-course northern blot analysis of rRNAs extracted from wild-type Salmonella expressing SehA . Cells were harvested at indicated times after L-arabinose induction.

Given that *Salmonella* Typhimurium has seven rRNA operons and the sequences surrounding SehA cleavage sites are conserved, SehA cleavage sites were detected in all seven rRNA operons (Figures 3C and Figure S6). Because SehA induces cleavage in the leader region of 16S rRNAs, one can expect that SehA induction accumulates short leader RNA fragments and pre-16S rRNA fragments containing 115 nt of the leader region. To test this, we designed DNA probes to detect rRNAs containing the leader regions (L1 and L2) and only mature rRNAs (R3 and R4). Northern blot analysis showed that SehA induction indeed accumulated short leader RNAs (detected by the L1 probe) and pre-16S RNAs (detected by the L2 probe) but not mature 16S rRNAs (detected by the R3 and R4 probes) (Figures 3D and 3E). As controls, the vector-expressing or non-inducing control did not increase leader fragments or pre-16S rRNAs (Figure 3D). These data indicate that SehA toxin induces cleavage of rRNAs and accumulates short leader RNAs and pre-16S rRNAs *in vivo*.

### The SehA toxin-induced cleavage site in the 16S rRNA leader coincides with the RNase III cleavage site

It turns out that SehA cleavage site at the -115 position in the 16S rRNA leader is identical to the RNase III cleavage site reported in *E. coli* (Bram et al., 1980; Young and Steitz, 1978)(Figure 4). Bacterial rRNA is transcribed as a single transcript and processed into mature 16S, 23S, and 5S rRNAs by multiple ribonucleases (Bechhofer and Deutscher, 2019; Court et al., 2013). Among these ribonucleases, RNase III is involved in the primary step in rRNA processing by cleaving at the double-stranded stem regions that base-pair between sequences 5′ and 3′ to rRNAs (Figure 4A). In the 16S rRNA region, RNase III cleavage generates a precursor of 16S rRNA containing an extra 115 nt and 21 nt at the 5′ and 3′ ends, respectively, which are further processed by RNase E and RNase G to generate mature 16S rRNAs (Bechhofer and Deutscher, 2019; Court et al., 2013).

**Figure 4.**
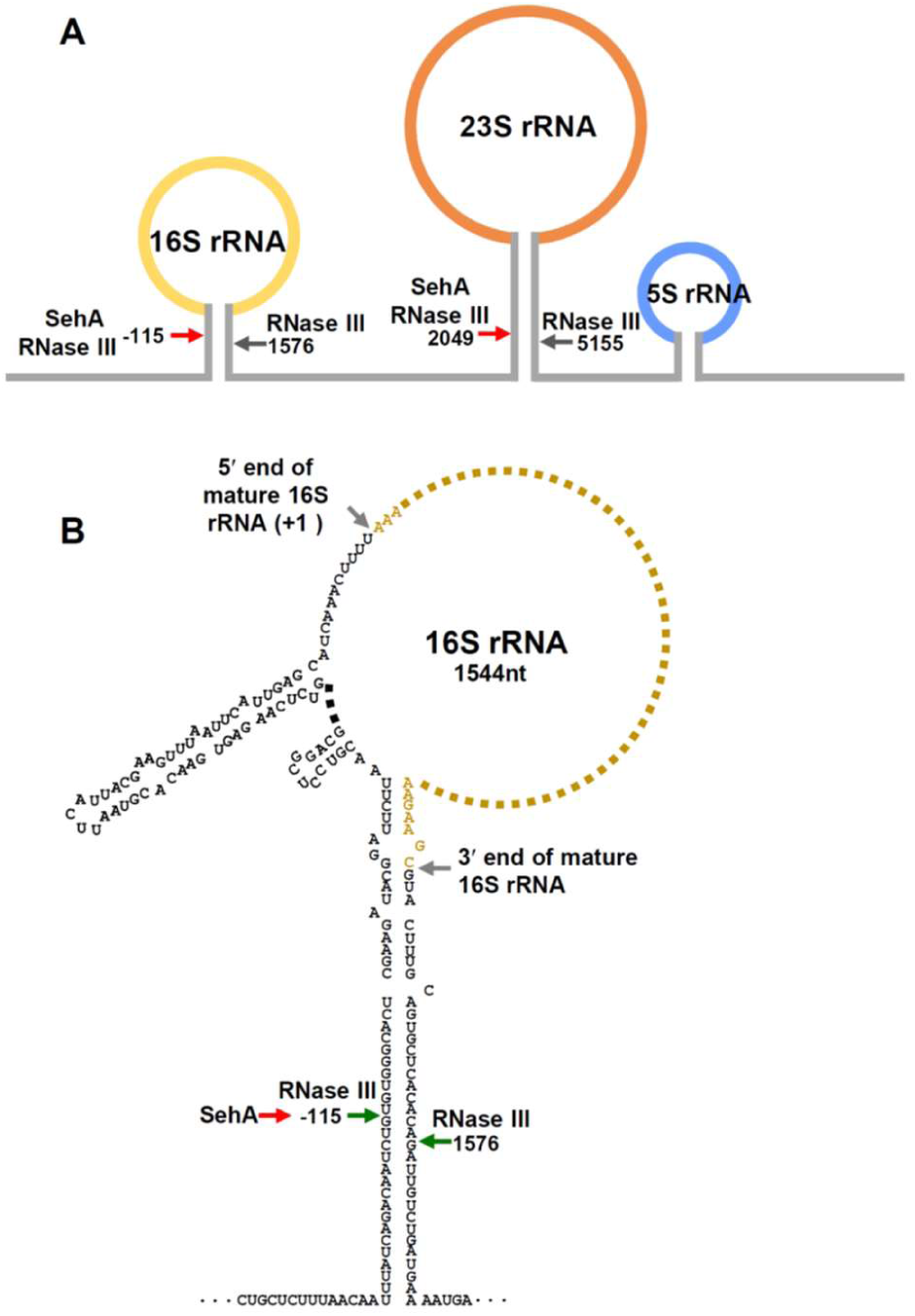
SehA toxin has an identical cleavage site with RNase III within the leader region of 16S rRNA. (A) Schematic view of Salmonella enterica 16S-23S-5S rRNA operon. The mature sequences of 16S, 23S, and 5S are indicated in yellow, orange, and blue, respectively. Black bold arrows indicate positions of RNase III cleavage sites, which correspond to the nucleotide positions relative to the first nucleotide (+1) of the 16S rRNA mature sequence. Red arrows correspond to SehA toxin cleavage sites, which were mapped in vivo. (B) Predicted secondary structure of the pre-16S rRNA. The mature 16S rRNA are shown in yellow and the 5’ and 3’ ends of 16S rRNA are indicated by grey arrows. The leader and spacer sequences are shown in black. Each number corresponds to the nucleotide position relative to the 5’ of the mature 16S rRNA (+1). Green arrows indicate RNase III cleavage sites and red arrow at position -115 indicates SehA cleavage site identical to one of the RNase III cleavages sites.

If SehA induces cleavage in the 16S rRNA leader at the same location as RNase III and accumulates 16S RNA precursors, we wondered what the consequence of this event would be. Since RNase III-mediated rRNA cleavage is an initial step of rRNA maturation and ribosome assembly, one might expect that SehA-induced accumulation of rRNA precursors and a decrease in mature rRNA indicate a failure or delay in further rRNA processing and ribosome assembly. To test this idea, we determined ribosome profiles prepared from *Salmonella* expressing SehA. *Salmonella* expressing the empty vector exhibited peaks (A_260_) corresponding to 30S, 50S, and 70S ribosome fractions (Figures 5A and 5D). However, SehA induction decreased the overall heights of all three peaks and showed indistinct boundaries between peaks (Figures 5B and 5E), suggesting that SehA toxin disrupts ribosome assembly by preventing rRNA maturation. As a control, *Salmonella* expressing a nonfunctional SehA variant with the Y53A substitution retained individual peaks of 30S, 50S, and 70S ribosome fractions (Figures 5C and 5F).

**Figure 5.**
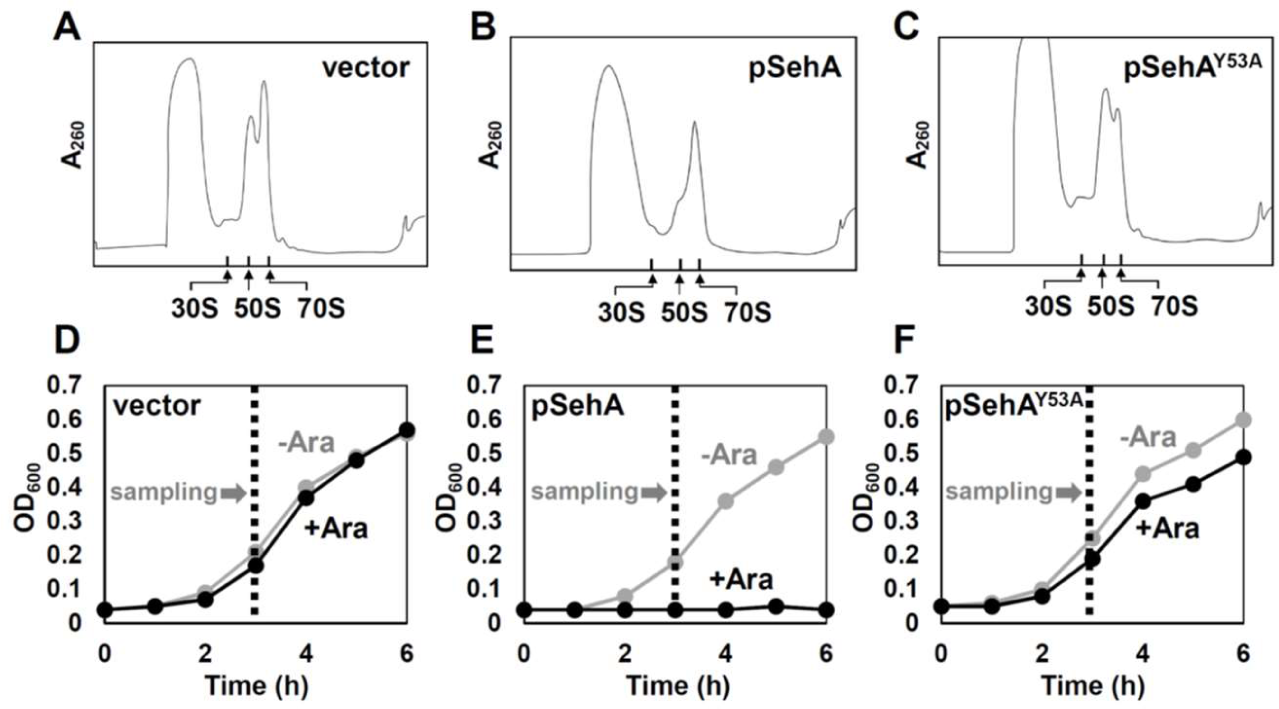
SehA toxin expression reduces 70S ribosome assembly. (A-C) Ribosome profile analyses of Salmonella strains expressing the empty vector (A), wild-type SehA toxin (B), or SehA Y53A variant grown in N-minimal medium containing 0.01 mM Mg^2+^ and 10 mM L-arabinose. Extracts from above strains were fractionated on 10-50% sucrose gradients and the absorbance at 260 nm was measured during fractionation. The peaks of 30S, 50S, and 70S are indicated by arrows. (D-F) Growth curves of the above strains grown in N-minimal medium containing 0.01 mM Mg^2+^ at 37°C for 6 h with shaking and measured absorbance at 600 nm every hour. Sampling time-points for ribosome profile analyses are indicated by dotted lines.

### SehA’s toxin activity is dependent on RNase III

Because SehA alone does not exhibit endoribonuclease activity *in vitro* (Figure S2) and also because *in vivo* SehA cleavage sites coincide with RNase III cleavage sites (Figure 4), we suspected that SehA’s toxin activity may require RNase III. To test this, we created a chromosomal deletion mutant of the *rnc* gene encoding RNase III. Similarly to what we observed previously (Figures 1B and 1C), SehA induction decreased growth compared to the non-inducing control or the empty vector expression (Figures 6A and 6B). Although it was reported that the *rnc* gene is dispensable in *E. coli* (Takiff et al., 1989), the *Salmonella rnc* deletion delayed growth compared to the wild-type (Figure 6C). Interestingly, SehA induction did not further reduce growth in the *rnc* mutant background. Instead, SehA expression slightly increased growth compared to the SehA-non-inducing control in the *rnc* background (Figure 6D). By contrast, SehA overexpression further decreased growth in the *rng* mutant, which encodes another rRNA processing ribonuclease, RNase G (Figures S7A and S7B). These data suggest that SehA exhibits its toxin activity in the presence of RNase III.

**Figure 6.**
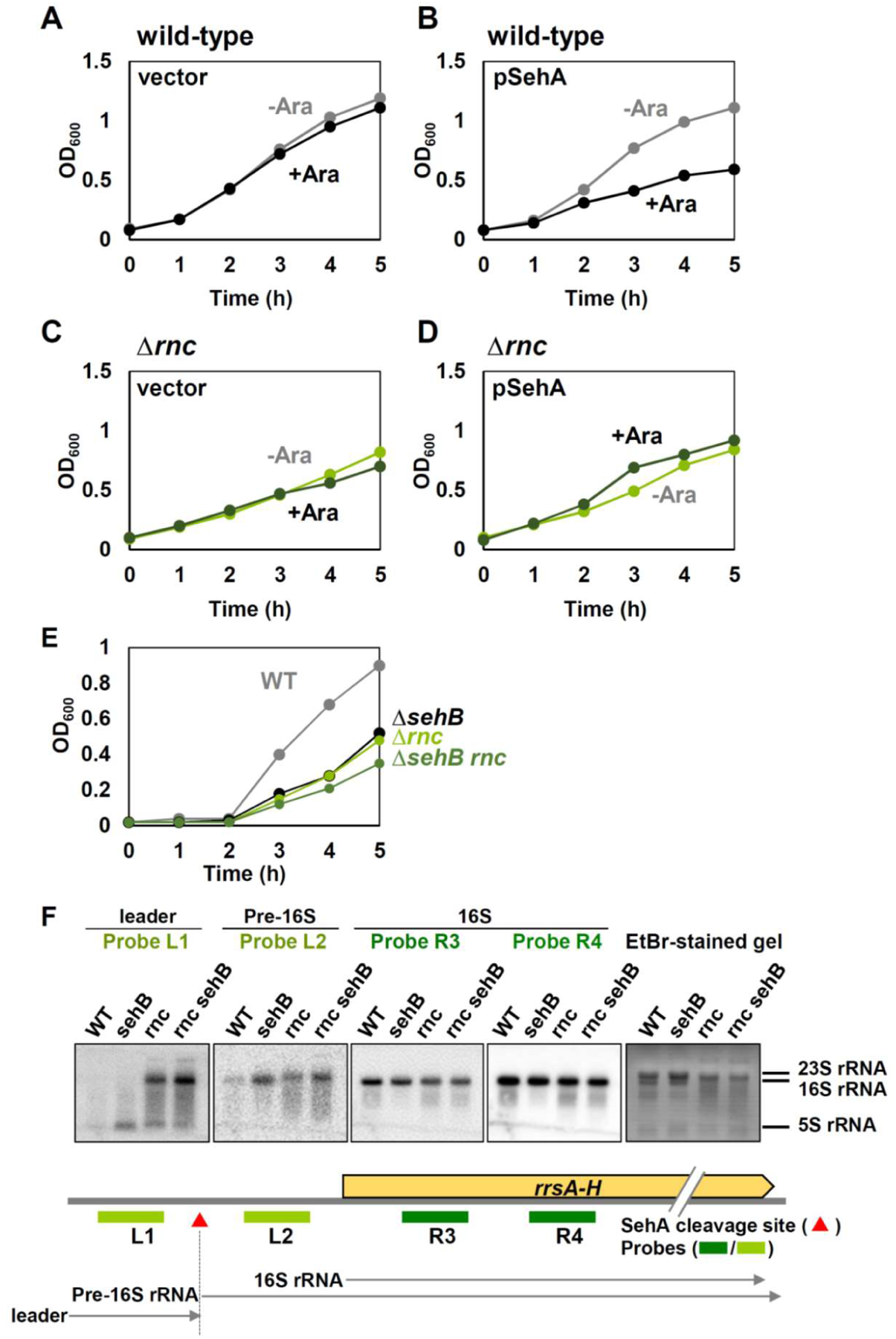
SehA toxin’s effect is RNase III-dependent. (A-D) Growth curves of the wild-type or rnc deletion Salmonella strains expressing the empty vector (A and C) or pSehA (B and D) in N-minimal medium containing 10 mM Mg^2+^ in the presence or absence of 1 mM L-arabinose. Bacteria were grown at 37°C for 5 h with shaking and measured absorbance at 600 nm every hour. (E-F) Chromosomal sehB deletion has a marginal effect on rnc mutant. (E) Growth curves of wild-type, the sehB deletion mutant (ΔsehB), rnc deletion mutant (Δrnc), and a strain deleted both the sehB and rnc genes (ΔsehB rnc) grown in N-minimal medium containing 10 mM Mg^2+^ at 37°C for 5 h with shaking and measured absorbance at OD600 every hour. (F) Northern blot analysis of rRNAs extracted from Salmonella strains listed above. Total RNAs were separated on a MOPS-agarose gel and stained with ethidium bromide (EtBr, far right). The gel was transferred on a membrane and then the membrane was hybridized with probes L1, L2, R3, and R4. The SehA cleavage sites are indicated as red tringles. The mature 16S rRNA and the precursors (leader and pre-16S) are indicated below.

To confirm SehA’s effect in the *rnc* background from its chromosomal location, we created a chromosomal mutant where both the *sehB* and *rnc* genes were deleted. In N-minimal media, the *sehB* deletion mutant showed a growth defect similar to the *rnc* single deletion mutant (Figure 6E), indicating that the effect of SehA expression from its chromosomal location by deleting *sehB* has a similar growth defect to the *rnc* deletion. When we combined *sehB* and *rnc* mutations, *sehB* deletion had a marginal effect on growth in the *rnc* deletion mutant (Figure 6E). Thus, SehA’s effect on growth in the *sehB* deletion mutant is largely dependent on RNase III. However, the *rnc* mutation did not alter mRNA expression behaviors of the *sehAB* genes (Figures S7C-S7E).

We also measured rRNA fragments generated in the *sehB* or *rnc* mutants using Northern blot analyses. Similarly to what we detected in the strain heterologously expressing SehA (Figure 3D), the *sehB* deletion accumulated leader rRNA fragments detected by the L1 probe (Figure 6F, probe L1). In addition, the *rnc* deletion exhibited less clear boundaries of leader rRNA fragments, reflecting inefficient rRNA processing due to the *rnc* deletion. Still, the *sehB* deletion in the *rnc* mutant did not increase rRNA leader cleavage fragments (Figure 6F, probe L1). Pre-16S rRNA fragments detected by the L2 probe showed patterns similar to those of leader rRNA fragments (Figure 6F, probe L2), indicating that SehA induces rRNA cleavage in an RNase III-dependent fashion. Levels of mature 16 rRNA fragments slightly decreased when the *sehB* gene was deleted, but the *sehB* deletion had no further decrease in mature rRNA levels of the *rnc* deletion mutant (Figure 6F, probes R3 and R4).

### SehA exerts its activity by binding to RNase III

We wondered how SehA exerts its toxin activity in an RNase III-dependent manner. First, we tested whether SehA can bind to double-stranded RNA similarly to RNase III. To test this, we incubated purified SehA proteins with increasing concentrations of *in vitro* synthesized double-stranded RNA templates that correspond to the stem region of the *rrsH* 16S rRNA leader. In agreement with the previous result (Figure S2), SehA does not slow down the mobility of *in vitro* synthesized 16S leader RNA (Figure S8), indicating that SehA does not directly bind to 16S rRNA *in vitro*.

Second, we tested whether SehA then interacts with RNase III (Figure S9A and S9B). When GFP-tagged SehA proteins were immobilized with beads, SehA proteins were co-eluted with His-tagged RNase III proteins (Figure S9C). Interestingly, SehA Y53A variant did not alter the elution profile (Figure S9D), suggesting that SehA toxin’s activity does not affect RNase III binding. In bacteria, RNase III contains a ribonuclease domain (RND) and a dsRNA-binding domain (dsRBD) (Figure S9A and S9B). To search for the minimal region required for SehA binding, we also tested with purified His-tagged dsRNA-binding domain (dsRBD) alone and found that both wild-type or Y53A variant SehA toxins were still co-eluted with the dsRBD domain (Figures S9E and S9F), indicating that SehA toxin binds to the dsRNA-binding domain of RNase III independently of the toxin’s activity.

Third, to confirm whether SehA exerts its toxin activity by binding to RNase III, we purified wild-type or Y53A-substituted SehA and RNase III proteins and reconstituted SehA-mediated rRNA cleavage *in vitro*. For this, 88 nt-long *in vitro* synthesized transcripts that retain the stem region with RNase III cleavage sites but remove most of the mature 16S rRNA were incubated with SehA and/or RNase III (Figures 7A and 7B). Similarly to Figure S2, SehA alone did not produce cleavage fragments when mixed with the 88-nt RNA substrate, while RNase III cleaves the RNA substrate, generating 22 nt, 35 nt, and 31 nt cleavage products (Figure S10). When incubated with different amounts of RNase III, RNase III alone cleaves the RNA substrates, generating cleavage products of 22 nt, 35 nt, and 31 nt at 4 pmol (Figures 7C and 7D). However, addition of SehA^WT^ proteins did not produce cleavage fragments at those concentration of RNase III, suggesting that SehA toxin interferes with the normal activity of RNase III (Figures 7C and 7D), probably by binding to the dsRBD of RNase III (Figure S9). As a control, the addition of SehA^Y53A^ did not interfere RNase III activity (Figures 7C and 7D).

**Figure 7.**
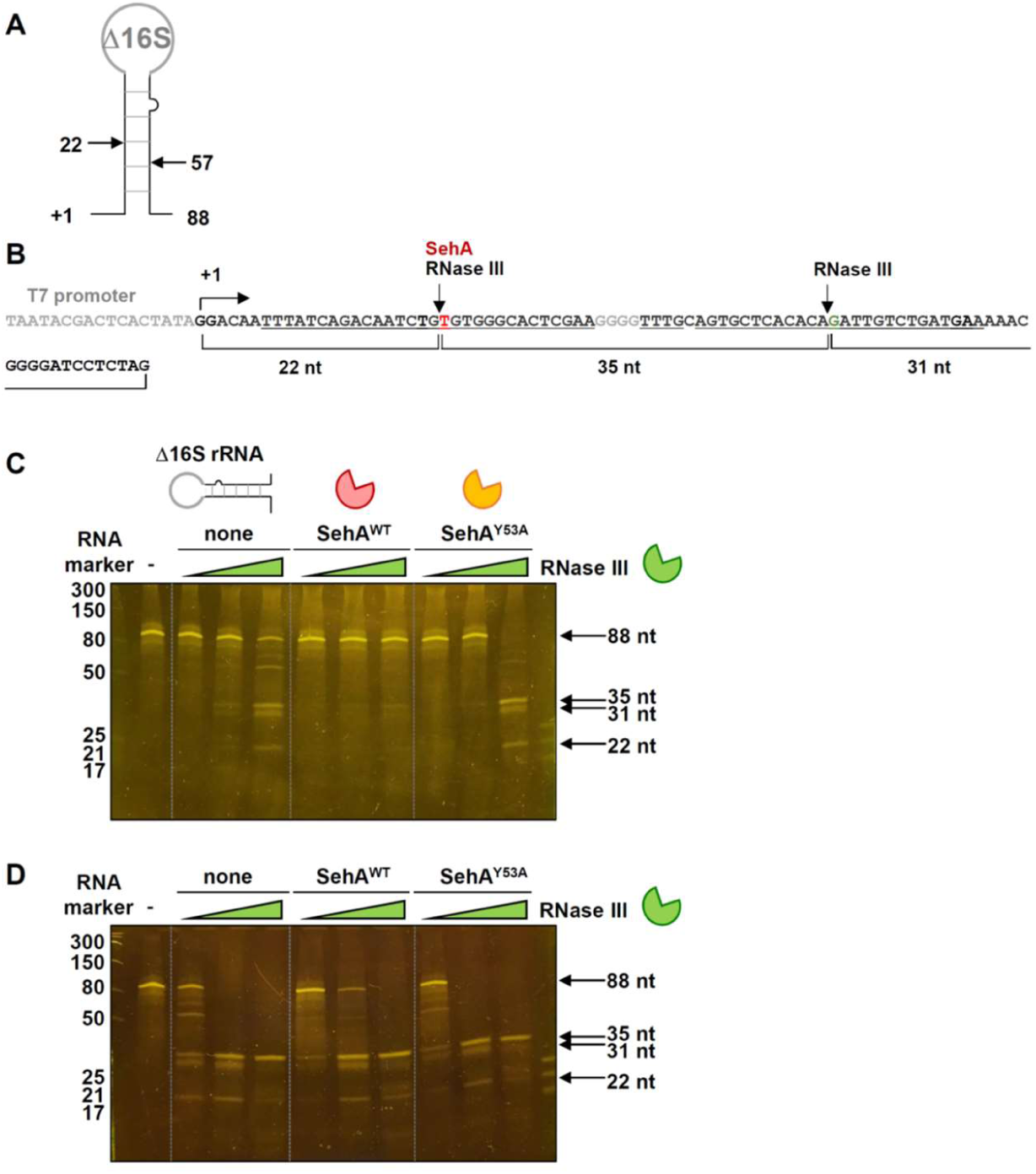
SehA toxin inhibits RNase III-mediated 16S rRNA cleavage in vitro. (A) Schematic diagram of in vitro synthesized 16S rRNA substrate. rRNA transcripts were synthesized in vitro, 88 nt RNA corresponding to sequence lacking most of the 16S rRNA mature sequence but retaining the stem structure flanking the 16S rRNA. (B) Sequence of DNA template for in vitro synthesized rRNA substrate. (C-D) In vitro RNase III cleavage assay of in vitro synthesized 16S rRNA substrate. The rRNA substrates were incubated for 10 min with the increasing amounts of Salmonella RNase III (C: 0.12, 0.25, and 1 pmol and D: 1, 4, and 16 pmol), either in the presence or absence of SehA^WT^ or SehA^Y53A^. Controls with RNA alone were included. The reaction products were separated on a 15% TBE-urea gel and visualized by SYBR gold staining.

Collectively, these results indicate that SehA toxin binds to RNase III via the dsRNA-binding domain (dsRBD) and inhibits RNase III from processing the initial step of rRNA maturation. SehA-mediated RNase III inhibition induces the accumulation of pre-rRNA fragments, presumably interfering with further processing by other RNases. This action results in reduction of mature rRNA production and ribosome assembly, leading to a slowdown in bacterial growth.

### SehA induces non-replicating *Salmonella* within macrophages in an RNase III-dependent manner

When we infected macrophage-like J774A.1 cell lines with wild-type *Salmonella*, the *sehA* gene was highly induced inside macrophages (∼370-fold at 6 hours post-infection; Figure S11A), suggesting that SehA toxin is required during infection. The *sehB* gene was also induced but at slightly lower levels (∼91-fold at 6 hours post-infection; Figure S11B), possibly because the *sehA* toxin gene precedes the *sehB* antitoxin gene.

We then wondered about the physiological significance of SehA-mediated RNase III inhibition during *Salmonella* infection. To explore this, we infected macrophage-like J774A.1 cell lines with *Salmonella* strains to measure *Salmonella* survival within macrophages. Similar to what we observed *in vitro* (Figures 1D and 1E), the *sehB* mutant decreased *Salmonella* replication within macrophages by ∼50% compared to wild-type *Salmonella* at 21 hours post-infection (Figures 8A and 8B). This indicates that, within macrophages, the *sehB* mutant *Salmonella* expresses SehA sufficiently to slow down growth inside the host. Interestingly, the *rnc* mutant decreased *Salmonella* survival similar to the *sehB* mutant, but *sehB* deletion in the *rnc* mutant background did not further decrease *Salmonella* survival as detected in the wild-type background (Figure 8A). This data suggests that the survival defect of the *sehB* mutant inside macrophages is dependent on RNase III.

**Figure 8.**
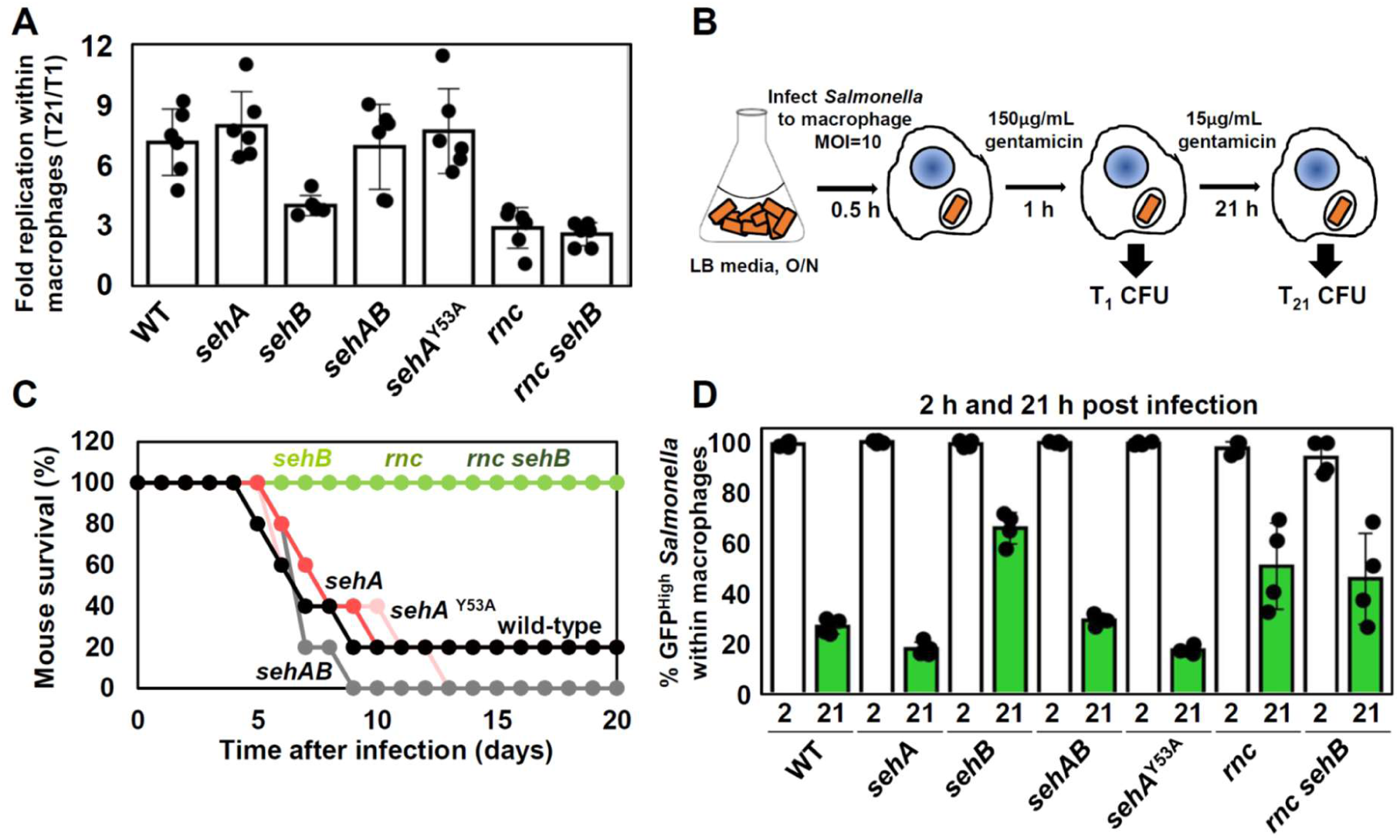
SehA induces non-replicating cells inside macrophages in an RNase III-dependent manner. (A) Survival inside J774A.1 macrophages of the Salmonella strains including wild-type, sehA, sehB, sehAB, rnc, and sehB rnc, or a strain with the sehA Y53A substitution at 21 h post infection (T21). Fold replication represents [number of bacteria at T21/number of bacteria at T1] as shown in (B). Shown are the means and SD from six independent infections. (B) Schematic diagram of macrophage survival assay. (C) Survival of C3H/HeN mice inoculated intraperitoneally with ∼10^3^ colony-forming units of the Salmonella strains listed above. (D) Fluorescence dilution assay of the Salmonella strains listed above inside J774 A. 1 macrophages. The percentage of cells expressing high levels of GFP (GFPhigh) was calculated from flow cytometric detection of mCherry and GFP fluorescence in Salmonella harboring pFCGi plasmid (n = 12,000 cells). Shown are the means and SD from four independent infections. *P < 0.05, n.s. not significant, two-tailed t-test. See also Figure S11.

Likewise, the *sehB* mutant as well as *rnc* and *sehB rnc* double mutants were completely avirulent when mice were inoculated with ∼10^3^ colony-forming units (CFUs) of those *Salmonella* strains intraperitoneally (Figure 8C). Thus, these data collectively indicate that SehA expression via *sehB* deletion decreases *Salmonella* survival within macrophages and attenuates virulence in mice.

However, in both cases, *sehA* deletion or a chromosomal mutant with the *sehA* Y53A substitution did not exhibit significant differences compared to wild-type (Figure 8A). Given that the toxin module in the toxin-antitoxin system is responsible for producing a fraction of non-replicating cells within the bacterial population and this can be undetectable by CFU counting (Gollan et al., 2019; Jurenas et al., 2022), we then measured to see whether SehA contributes to the formation of non-replicating *Salmonella* within macrophages at the single-cell level. To explore this, we utilized *Salmonella* harboring a plasmid expressing GFP from an arabinose-inducible promoter and mCherry from a constitutive promoter (Helaine et al., 2014)(Figure S11C). After GFP-expressing *Salmonella* infected macrophages without arabinose, we measured the percentage of *Salmonella* retaining high GFP levels (GFP_high_) as non-replicating *Salmonella*; otherwise, GFP levels would be diluted as *Salmonella* replicates within macrophages (Figures S11D and S11E). Wild-type *Salmonella* retained 26.3±3.0% of GFP-high cells inside macrophages at 21 hours post-infection (Figure 8D). In this case, SehA is responsible for maintaining GFP_high_ cells because *sehA* deletion or *sehA* Y53A substitution significantly decreased percentages of non-replicating cells (Figure 8D)(18.4±3.3% and 18.8±1.7%, respectively). By contrast, SehA expression by *sehB* deletion greatly increased GFP_high_ cells up to 65.1±6.4%. Combining both *sehA* and *sehB* deletion recovered GFP_high_ cells similar to wild-type (Figure 8D). Likewise, *rnc* deletion increased the percentage of GFP_high_ cells similar to those in the *sehB* mutant (Figure 8D). However, *rnc sehB* double deletion did not further increase the percentage of GFP_high_ cells (Figure 8D), supporting that SehA maintains non-replicating *Salmonella* during host infection by inhibiting RNase III. As controls, the percentages of GFP_high_ cells in all tested strains were similar at 2 hours post-infection (Figure 8D).

## Discussion

In this study, we have established that SehA, a *Salmonella* HigB-like toxin, inhibits *Salmonella* growth and induces the formation of non-growing *Salmonella* within macrophages. SehA toxin exerts its regulatory effect by binding to RNase III via its dsRNA-binding domain (dsRBD) and inhibits RNase III-mediated rRNA processing. Because RNase III catalyzes the initial step of rRNA processing, SehA-mediated rRNA processing inhibition results in a delay in ribosome assembly (Figure 9). In bacteria, the number of ribosomes directly correlates with growth rate under given conditions (Lindahl and Zengel, 1986; Nomura et al., 1984). Thus, the delay in ribosome assembly slows down growth and increases the fraction of non-replicating *Salmonella* during infection.

**Figure 9.**
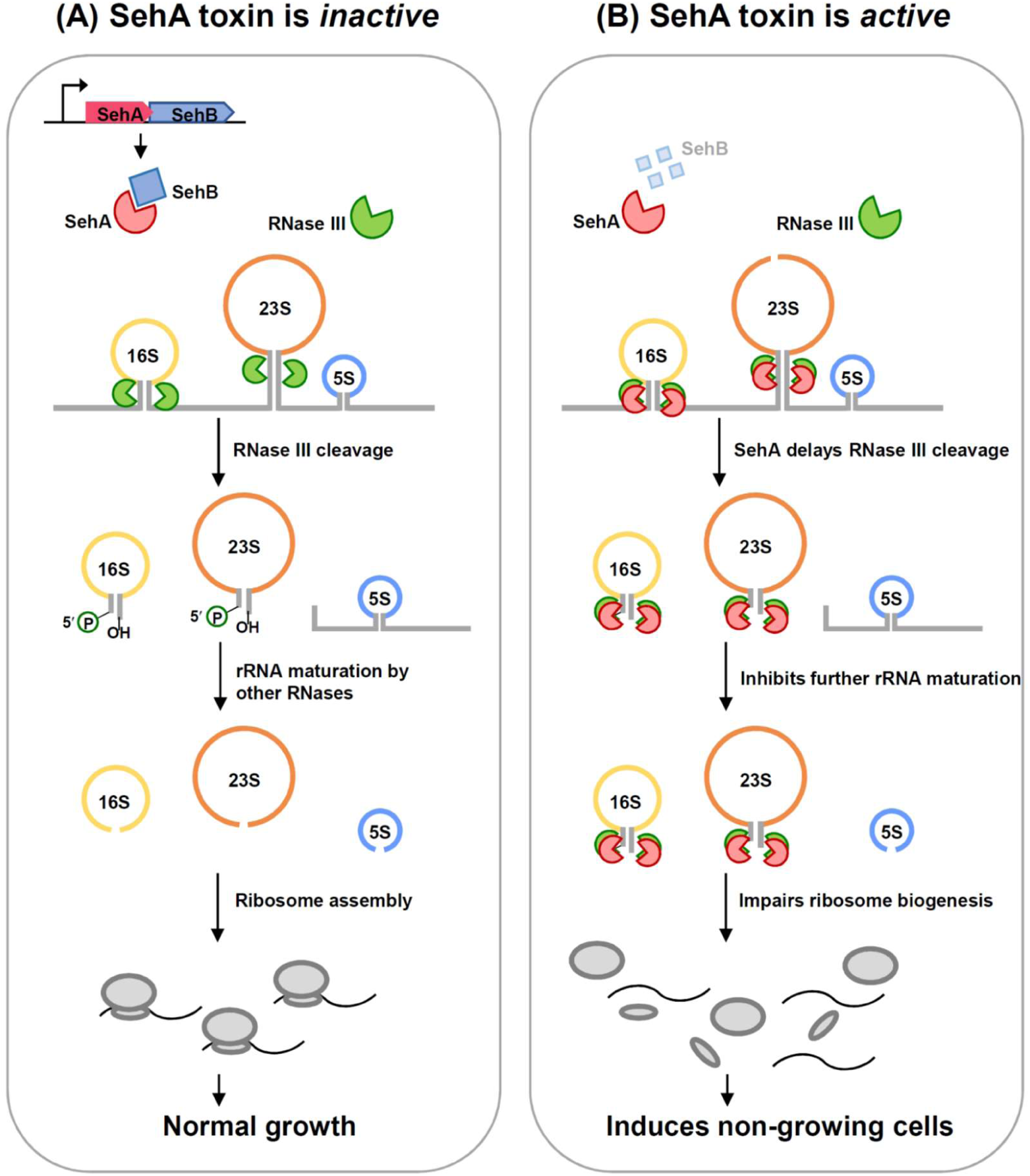
SehA toxin-mediated inhibition of rRNA maturation promotes persister cells inside macrophages. In the presence of SehB antitoxin, SehA toxin binds to SehB antitoxin. Meanwhile, RNase III processes newly transcribed rRNA transcript and facilitates maturation of 16S, 23S, and 5S rRNAs, which are assembled into 70S ribosome. In the absence of SehB, SehA toxin is released and binds to RNase III, which inhibits RNase III-mediated rRNA processing in the leader regions of 16S and 23S rRNAs. Because SehA delays RNase III-mediated rRNA processing, which appear to inhibit further maturation of rRNA by other ribonucleases. This prevents from 70S ribosome assembly that gives rise to induce non-growing cells in antibiotic treatment or during macrophage infection.

Overall, SehA toxin is homologous to HigB toxins from *E. coli* and *P. vulgaris* (Hurley and Woychik, 2009; Yang et al., 2016). The HigB toxins belong to the RelE-type superfamily of toxins, which are predominantly ribosome-dependent mRNA endoribonuclease (Han and Lee, 2020). Although RelE-type toxins possess a wide range of mRNA codon specificity depending on their structural features to interact with mRNA and ribosomes (Han and Lee, 2020), they mainly cleave between the 2^nd^ and 3^rd^ nucleotides in the ribosomal A-site. Similarly, *P. vulgaris* HigB is also known to cleave ribosome-bound mRNAs in an adenosine-rich codon-dependent manner (Hurley and Woychik, 2009). Based on the structural similarities, SehA was initially expected to be ribosome-dependent endoribonuclease toxin.

Strikingly, however, RNA sequencing analysis to search for 5′ hydroxylated (5′ OH-) or 5′ monophosphorylated (5′ monoP-) RNA ends generated upon SehA induction revealed that i) SehA-mediated RNA fragments were located in non-coding RNAs, mostly in the leader regions of rRNA operons, and ii) there were no significant differences between the sequencing results of RNA fragments with both 5′ OH- and 5′ monoP-ends and those with only 5′ monoP-ends. This suggests that SehA’s molecular activity can differ from other RelE/HigB-type toxins and that most SehA-induced RNA fragments contain 5′ monoP-ends (Figure 3). The fact that SehA-induced 5′ monoP-RNA ends in the rRNA operon coincide with RNase III processing sites (Figure 4) and that SehA alone does not cleave RNA (Figure S3) led us to discover that SehA toxin itself is not an endoribonuclease but acts on RNase III to inhibit *Salmonella* growth.

Therefore, despite sharing structural similarities, these similarities do not necessarily imply functional similarities. A similar situation can be found in Kid-like family toxins including Kid, MazF, and CcdB toxins, which share structural similarities but exhibit different functions (Kid and MazF are endoribonucleases whereas CcdB is a gyrase inhibitor.)(Hargreaves et al., 2002; Kamada et al., 2003; Loris et al., 1999).

Pre-16S rRNA harboring 115 nt of the leader region is not normally detected in wild-type (only 1∼2% of rRNA), as it is rapidly processed further into mature 16S rRNA by other RNases (King and Schlessinger, 1983; Srivastava and Schlessinger, 1990). Thus, the accumulation of pre-16S rRNA upon SehA induction reflects a situation where normal rRNA processing is likely disrupted (King and Schlessinger, 1983; Srivastava and Schlessinger, 1990). Similarly, a strain lacking RNase III accumulated pre-16S rRNA as well as a longer precursor containing 23S rRNA, and *sehB* deletion in the *rnc* background did not further increase pre-16S rRNA (Figure 6), suggesting that SehA-mediated accumulation of pre-16S rRNA can be due to SehA’s inhibitory action on RNase III. Indeed, this is further supported by an experiment in which purified SehA directly inhibits RNase III-mediated cleavage of the 16S RNA leader stem *in vitro* (Figure 7).

It is interesting to note that SehA incubation inhibits RNase III-mediated 16S RNA processing *in vitro*, thus generating fewer cleavage products of 16S rRNA leader region. However, *in vivo* SehA induction accumulated more pre-16S rRNA fragments and decreased mature 16S RNA compared to the non-induced control. We interpreted this difference as a consequence of disruption of rRNA processing because the entire rRNA maturation process is tightly coupled/coordinated with subsequent other RNases-mediated RNA maturation and ribosome assembly (King and Schlessinger, 1983; Srivastava and Schlessinger, 1990). Therefore, SehA-mediated inhibition or delay in the initial rRNA processing step by RNase III can cause uncoupling of this event from the remaining maturation steps, thereby accumulating yet-to-be-processed pre-16S RNA. This uncoupling appears to decrease ribosome assembly (Figures 5 and 9), resulting in a growth defect. Although SehA itself does cleaves mRNA as an endoribonuclease, it directly targets RNase III and thus inhibits RNase III-mediated rRNA processing and ribosome assembly. Therefore, it also constitutes a translation-inhibiting toxin, which is most predominant among toxin-antitoxin systems (Harms et al., 2018).

SehA was predicted to be an endoribonuclease based on homology but did not exhibit endoribonuclease activity. Instead, it directly binds to the dsRNA-binding domain (dsRBD) of RNase III and inhibits RNase III-mediated rRNA processing during *Salmonella* infection by an as-yet-unclear mechanism. Previously, *E. coli* YmdB protein was also reported to inhibit RNase III activity (Kim et al., 2008). Similarly to SehA toxin, YmdB directly binds to RNase III and functions as a *trans*-acting negative regulator of RNase III activity in response to cold-shock stress. However, unlike SehA, YmdB binds to the RNase/dimerization domain instead of the dsRNA-binding domain and inhibits the dimerization and activation of RNase III (Kim et al., 2008). Moreover, YmdB is predicted to be an O-acetyl-ADP-ribose deacetylase, the contribution of which to the inhibition of RNase III has not been elucidated.

A previous crystallographic study combined with modeling proposed that RNase III binds to dsRNA as a dimer. In each monomer, the N-terminal catalysis/dimerization domain (RND) and C-terminal dsRNA-binding domain (dsRBD) contact dsRNA at four RNA-binding motifs (4RBMs; 3-4 RBMs in the RND and 1-2 RBMs in the dsRBD) (Court et al., 2013; Gan et al., 2006). Because it is still unclear how both SehA or YmdB binding to the RNase III dsRBD or RND affects dsRNA cleavage, future structural studies with RNase III and its interacting proteins like SehA could provide insights for designing inhibitors that effectively target RNase III. Given that specific inhibitors for RNase III are currently unavailable and an engineered version of RNase III can be used to generate siRNA for gene silencing in mammalian cells (Xiao et al., 2009), these insights can be useful in many areas of research.

## MATERIALS AND METHODS

### Bacterial strains, plasmids, oligodeoxynucleotides, and growth conditions

Bacterial strains and plasmids used in this study are listed in Table S3. All *S. enterica* serovar Typhimurium strains were derived from the wild-type strain 14028s (Fields et al., 1986) and were constructed by one-step gene inactivation method (Datsenko and Wanner, 2000) and/or P22-mediated transduction as previously described (Davis et al., 1980). DNA oligonucleotides are listed in Table S2. Bacteria were grown at 37°C in Luria-Bertani (LB) broth and N-minimal media (Snavely et al., 1991) supplemented with 0.1% casamino acids, 38 mM glycerol, and the indicated concentrations of MgCl_2_. *E. coli* DH5α was used as the host for preparing plasmid DNA. Ampicillin was used at 50 µg mL^-1^, chloramphenicol at 20 µg mL^-1^, kanamycin at 50 µg mL^-1^, tetracycline at 10 µg mL^-1^, and fusaric acid (Maloy and Nunn, 1981) at 12 µg mL^-1^. IPTG (isopropyl β-D-1-thiogalactopyranoside) was used at 0.25 mM, L-arabinose at 0.2% (w/v) and X-Gal (5-bromo-4-chloro-3-indolyl β-D-galactopyranoside) at 80 µM.

### Protein structure modeling and protein docking modeling

We used Protein Homology/analogy Recognition Engine V 2.0 (Phyre2)(Kelley and Sternberg, 2009) to model structures of the SehA and RNase III proteins from *Salmonella enterica* serovar Typhimurium 14028s. The structure of the SehA protein was modelled based on the structure of mRNA interferase HigB from *E. coli* (PDB: 5ifg) and the structure of the RNase III protein was modelled based on the structure of RNase III from *E. coli* (PDB: 7R97). Then, we used ClusPro webserver to investigate a potential interaction between *Salmonella* SehA and RNase III proteins (Comeau et al., 2004).

### Quantitative real-time polymerase chain reaction (qRT-PCR)

Total RNA was isolated using RNeasy Kit (Qiagen), according to the manufacturer’s instructions. The purified RNA was quantified using a Nanodrop machine (ThermoFisher). cDNA was synthesized using PrimeScript RT reagent Kit (TaKaRa). The mRNA levels of the *sehA, sehB,* and *rnc* genes were measured by quantifying the cDNA using PowerUP^TM^ SYBR Green Master Mix (Applied Biosystems, A25742) and the appropriate primers (KHQ045/KHQ046 for the *sehA* gene, KHQ047/KHQ048 for the *sehB* gene, and KHQ143/KHQ144 for the *rnc* gene) and monitored using a StepOnePlus Real-Time PCR system (Applied Biosystems). The mRNA levels of each target gene were calculated using a standard curve of 14028s genomic DNA with known concentration and data were normalized to the levels of 16S ribosomal RNA amplified with primers 6970 and 6971.

### Protein expression and purification of His-tagged SehA, SehB, and RNase III proteins

BL21(DE3) cells harboring pRSF-sfGFP-*sehA*, pRSF-sfGFP-*sehB*, pET-His-FP-*rnc*, pET-His-FP-*rnc* RNase domain (1-128 aa), and pET-His-FP-*rnc* dsRNA binding domain (155-227 aa) were grown in 50 mL LB media at 37°C. When OD_600_ of the culture was reached to 0.5, IPTG was added to a final concentration of 1 mM and the bacteria were grown for 12 h at 25°C. Induced cultures were harvested, washed in 4 mL of TBS (Tris-buffered Saline) buffer, and lysed by sonication (Branson, digital sonifier 450 #60662). One milliliter of the cleared lysates was aliquoted for input samples, and the remaining lysates were incubated with 1 mL of Ni-NTA agarose (Qiagen, 1018244) on nutator (Daeiltech, PM6249) for 1 h. After settling on the column under gravity, His-tagged SehA, SehB, and RNase III proteins were purified according to the manufacturer’s instructions with modifications. Protein-bound resins were washed 10 volumes of wash buffer (50 mM Tris-HCl (pH 7.4), 150 mM NaCl, 0.5 mM EDTA, and 20 mM imidazole) and eluted with gradient elution buffer (50 mM Tris-HCl (pH7.4), 150 mM NaCl, 0.5 mM EDTA) with increasing concentrations of imidazole (50 mM, 100 mM, 200 mM, and 300 mM, respectively). Purified His-tagged proteins were concentrated and desalted using Amicon centrifugal filters (Millipore, UFC500396) in TBS buffer and stored in storage buffer (30 mM Tris–HCl, 0.5 M NaCl, 0.5 mM EDTA, 0.5 mM DTT, and 50% glycerol) at -80°C.

### In vitro RNA cleavage assay

*Salmonella* SehA and SehB proteins were purified from BL21 cells as described above. 10 µg of total RNAs were incubated in 50 µL of TBS (Tris-buffered Saline) at 25°C with 60 pmol of purified SehA proteins alone or SehA proteins preincubated with SehB proteins in a 1:1 molar ratio. At indicated times, 10 µL aliquots of the reaction were stopped by adding RNA sample loading buffer (R4268, Sigma). After incubation at 65°C for 2 min, the samples were resolved on a 1.5% MOPS-buffered formaldehyde-denaturing agarose gel for 30 min at 80 V (PowerPac^TM^ basic power supply, Bio-Rad) and visualized by gel imaging system (FluoroBox, CELLGENTEK).

### Pull-down assay between SehA toxin and RNase III proteins

The interaction between SehA toxin and RNase III protein were investigated using purified SehA, SehA ^Tyr 53 Ala^, RNase III, RNase III (dsRNA binding domain), and RNase III (RNase domain) proteins. Equimolar amounts of proteins were mixed with 25 µL of nickel beads in a total volume of 500 µL. The reactions were incubated at 4°C for 1 h on nutator and then the protein-bound beads were spin down by centrifugation and kept the supernatants for flow-through. The beads were washed with TBS washing buffer (20 mM Tris HCl pH 7.4, 150 mM NaCl) three times, and then the bound proteins were eluted in SDS sample buffer. The eluates were resolved on 12% SDS-polyacrylamide gels and stained with Coomassie dye staining or transferred to nitrocellulose membrane, and analyzed by Western blot using anti-His (1:5000 dilution, Rockland, 600-401-382) and anti-GFP (1:1000 dilution, Roche, 11814460001) antibodies for overnight. The blots were washed and hybridized with anti-rabbit IgG horseradish peroxidase-linked antibodies (1:10000 dilution, ThermoFisher, 31460) and anti-mouse IgG HRP-linked whole Ab (1:10000 dilution, Amersham, NA931) for 1 h and detected using SuperSignal West Femto Maximum Sensitivity Substrate (ThermoFisher).

### mRNA sequencing and analysis

Wild-type and *sehB* deletion *Salmonella* were grown overnight in N-minimal medium containing 10 mM Mg^2+^. A 1/100 dilution of the overnight culture was used to inoculate 10 mL of the same medium with 0.01 mM Mg^2+^, and grown for 4 h. Bacteria were harvested and RNA was isolated for further analysis. Total RNA was prepared by using RNeasy mini kit (Qiagen) and the integrity of the RNA samples was measured by using BioAnalyzer 2100 (Agilent Technologies). The samples with an RNA integrity number of over 8.0 were used for the next step. For ribosomal RNA depletion, 5 μg of the total RNA was processed by Ribo-Zero rRNA Removal Kit (Bacteria) (Illumina, MRZMB126). Sequencing libraries for RNA-Seq were constructed using NEBNext Ultra II Directional RNA Library Prep Kit for Illumina (NEB, E7760S), following the manufacturer’s instructions. Final sequencing libraries were quantified by PicoGreen assay (Life Technologies) and visualized using BioAnalyzer 2100. Sequencing was performed using a NextSeq 500 instrument (Illumina), following the manufacturer’s protocol, which generated 76 bp paired-end reads. The sequencing adapter removal and quality-based trimming for the raw data were performed using Trimmomatic v. 0.36 (Bolger et al., 2014). Cleaned reads were mapped to the reference genome (NCBI assembly; GCF_000022165.1) using bowtie2 with a default parameter (Langmead and Salzberg, 2012). For counting the reads mapped to each CDS (coding sequence), featureCounts was used (Liao et al., 2013). Finally, the count from each CDS was normalized using DeSeq2 package (Lee et al., 2024; Love et al., 2014). Processed data were deposited in the Gene Expression Omnibus (GEO) database with accession number GSE269990.

### 5’-ends identification of mono-phosphorylated RNA

5’-ends of hydroxylated or mono-phosphorylated RNA were identified using modified EMOTE protocol (Redder, 2015). Briefly, 1 µg of total RNA was processed by Ribo-Zero Magnetic Kit (Bacteria) (#MRZB12424, Illumina, USA). Resulting rRNA-depleted mRNA was phosphorylated by T4 polynucleotide kinase (#M0201, NEB, USA). The reaction was carried out at 37°C for 30 min in a total volume of 20 µL and heat-inactivated at 65°C for 30 min. Phosphorylated RNA was incubated with 4 µL of synthesized RNA oligo Rp6 (100 pmol) at 70°C for 5 min and cooled down. Rp6 was ligated to mono-phosphorylated transcripts at 37°C for 30 min by adding T4 RNA ligase 1 (#M0204, NEB, USA) with the buffer. After the reaction, 1 µL of 10 mM ATP was added to the sample and incubated at 16°C for overnight to complete ligation. RNA-oligo ligated transcripts were purified in 12 µL volume by AMPure XP (#A63881, Beckman Coulter, USA). Purified RNA was reverse-transcribed using SuperScript II (#18064-014, Invitrogen, USA) and the DROAA primer, which contains sequencing barcode, in a total volume of 20 µL reaction by manufacture’s protocol. Resulting cDNA was proceeded to PCR amplification using linker-(D6X) and DROAA-specific (B-PE-PCR20) primers. After 5 cycles, A-PE-PCR10 primer was added and continue 25 cycles to amplify PCR products between 300 and 800 bp.

### Sequencing and mapping with 5’-ends of mono-phosphorylated RNA

Generated library was sequenced by MiSeq platform (Illumina, USA) with 250 bp single end mode. The raw data was adapter- and quality-trimmed by using Trimmomatic v.0.36 (Bolger et al., 2014). Cleaned reads were mapped to the reference genome (*S. enterica* subsp. *enterica* serovar Typhimurium str. 14028s, PRJNA33067; NC_016856.1 and NC_016855.1 for pSLT1 plasmid) by hisat2 (Kim et al., 2019) with modified parameters (-k 1 --no-spliced-alignment). The mapping result was converted to BAM format and split to have a stranded specificity with SAM flag information. Resulted data was further modified to report the depth of 5′ site in the mapped read position using genomeCoverageBed (Quinlan and Hall, 2010) with changed parameters (-5 -dz). The reads were quantified per base using Integrated Genome Viewer (IGV) tools. Raw data was deposited in NCBI GEO database (accession number GSE269989).

### Plasmid construction

The plasmids pBAD33-*sehA*, pBAD33-*sehA* ^Tyr 53 Ala^, and pBAD33-*rnc* were constructed as follows: PCR fragments corresponding to the *sehA* and *rnc* genes were generated by PCR with the primers KHU859/KHU860 (for *sehA*) and KU280/ KU281 (for *rnc*) using 14028s genomic DNA as a template. A DNA fragment for the *sehA ^Tyr 53 Ala^* gene was generated by PCR with the primers KHU859/KHU860 using EN1072 genomic DNA as a template. The amplified DNA fragments were digested with XbaI and HindIII and cloned into pBAD33 digested with the same enzymes. The sequences of the resulting constructs were verified by DNA sequencing.

For protein purification, the plasmid pRSF-sfGFP-*sehA*, pRSF-sfGFP-*sehA* ^Tyr 53 Ala^, pRSF-sfGFP-*sehB*, pET-His-FP-*rnc,* pET-His-FP-*rnc* RNase domain (1-128 aa), and pET-His-FP-*rnc* dsRNA binding domain (155-227 aa) were constructed as follows: PCR fragments corresponding to the *sehA*, *sehB*, and *rnc* genes were generated by PCR with the primers KHU897/KHU898 (for *sehA*), KHU899/KHU900 (for *sehB*), KU285/KU287 (for *rnc*), KU285/KU552 (for *rnc* RNase domain), and KU551/KU287 (for *rnc* dsRNA binding domain) using 14028s genomic DNA as a template. For the *sehA* ^Tyr 53 Ala^ gene, a DNA fragment was generated by PCR with the primers KHU897 and KHU898 (for *sehA* ^Tyr 53 Ala^) using EN1072 genomic DNA as a template. For *sehA*, *sehB*, *sehA* ^Tyr 53 Ala^, and *rnc*, the amplified DNA fragments were digested with BamHI and XhoI and cloned into pRSF-sfGFP or pET-His-FP digested with the same enzymes. The sequences of the resulting constructs were verified by DNA sequencing.

### Construction of strains with chromosomal deletions of the *sehA*, *sehB*, or *rnc* **genes**

*Salmonella* strains deleted for *sehA, sehB,* or *rnc* genes were generated by the one-step gene inactivation method (Datsenko and Wanner, 2000). Cm^R^ cassettes for the *sehA, sehB, sehAB,* or *rnc* genes were PCR amplified from plasmid pKD3 using primers KHU634/KHU635 (for *sehA*), KHU944/KHU637 (for *sehB*), KHU634/KHU637 (for *sehAB*), or KU288/KU289 (for *rnc*) and the resulting PCR products were integrated into the 14028s chromosome to generate EN1011 (*sehA*::Cm^R^), EN1098 (*sehB*::Cm^R^), EN1013 (*sehAB*::Cm^R^), and SM368 (*rnc*::Cm^R^), respectively. The *sehA* (EN1019)*, sehB* (EN1108)*, sehAB* (EN1021), and *rnc* (SM371) strains were generated by removing Cm^R^ cassettes from EN1011, EN1098, EN1021, and SM368 via plasmid pCP20 as described (Datsenko and Wanner, 2000). P22 phage lysates grown in EN1098 (*sehB*::Cm^R^) strain were used to transduce SM371 (*rnc*) *Salmonella* selecting for chloramphenicol resistance to generate SM376 (*sehB*::Cm^R^ *rnc*). The *sehB rnc* (SM383) was generated by removing the Cm^R^ cassette from SM376 via plasmid pCP20 (Datsenko and Wanner, 2000).

### Construction of strains with chromosomal substitutions in the *sehA* gene

To generate strains with chromosomal mutations in the *sehA* coding region, we used the fusaric acid-based counter selection method as described previously (Lee and Groisman, 2010). First, we introduced a Tet^R^ cassette in the coding region of the *sehA* gene as follows: we generated a PCR product harboring the *tetRA* gene using primers KHU694/KHU695 and MS7953s chromosomal DNA as a template. The PCR product was purified using a QIAquick PCR purification kit (QIAGEN) and used to electroporate the 14028s chromosome containing plasmid pKD46 (Datsenko and Wanner, 2000). The resulting *sehA*::Tet^R^ (ENC1050) strain containing plasmid pKD46 were kept at 30°C. Then, we replaced the *tetRA* cassettes by preparing DNA fragments carrying various nucleotide substitutions in the *sehA* gene prepared by a two-step PCR process. For the first PCR reaction, we used two primer pairs KHU654/KHU753 and KHU752/KHU700 (for 53^rd^ tyrosine codon to alanine), KHU654/KHU755 and KHU754/KHU700 (for 61^st^ aspartate codon to alanine), KHU654/KHU691 and KHU690/KHU700 (for 88^th^ histidine codon to alanine), KHU654/KHU693 and KHU692/KHU700 (for 91^st^ tyrosine codon to alanine), KHU654/KHU714 and KHU713/KHU700 (for 93^rd^ lysine codon to alanine), KHU654/KHU716 and KHU715/KHU700 (for 95^th^ threonine codon to alanine), KHU654/KHU718 and KHU717/KHU700 (for 97^th^ tyrosine codon to alanine), and KHU654/KHU720 and KHU719/KHU700 (for 98^th^ tyrosine codon to alanine), and 14028s genomic DNA as a template. For the second PCR reaction, we mixed the two PCR products from the first PCR reaction as templates and amplified DNA fragments using primers KHU654 and KHU700. The resulting PCR products were purified and integrated into the EN1050 (*sehA*::Tet^R^) chromosome harboring pKD46 and selected against Tet^R^ with media containing fusaric acid to generate EN1072 (*sehA* ^Tyr 53 Ala^), EN1076 (*sehA* ^Asp 61 Ala^), EN1045 (*sehA* ^His 88 Ala^), EN1047 (*sehA* ^Tyr 91 Ala^), EN1078 (*sehA*^Lys 93 Ala^), EN1081 (*sehA* ^Thr 95 Ala^), EN1063 (*sehA* ^Tyr 97 Ala^), and EN1084 (*sehA* ^Tyr 98 Ala^), tetracycline-sensitive, ampicillin-sensitive (Tet^S^ Amp^S^) chromosomal mutants, respectively. The presence of the expected nucleotide substitutions was verified by DNA sequencing.

EN1099 (*sehA* ^Tyr 53 Ala^, *sehB*::Cm^R^), EN1100 (*sehA* ^Asp 61 Ala^, *sehB*::Cm^R^), EN1101 (*sehA* ^His 88 Ala^, *sehB*::Cm^R^), EN1102 (*sehA* ^Tyr 91 Ala^, *sehB*::Cm^R^), EN1103 (*sehA* ^Lys 93 Ala^, *sehB*::Cm^R^), EN1104 (*sehA* ^Thr 95 Ala^, *sehB*::Cm^R^), EN1105 (*sehA* ^Tyr 97 Ala^, *sehB*::Cm^R^), and EN1106 (*sehA* ^Tyr 98 Ala^, *sehB*::Cm^R^) mutants were generated by introducing Cm^R^ cassettes in the *sehB* gene of the above listed strains via one-step gene inactivation. Then, the EN1109 (*sehA* ^Tyr 53 Ala^, *sehB*), EN1110 (*sehA* ^Asp 61 Ala^ , *sehB*), EN1111 (*sehA* ^His 88 Ala^ , *sehB*), EN1112 (*sehA* ^Tyr 91 Ala^ , *sehB*), EN1113 (*sehA* ^Lys 93 Ala^, *sehB*), EN1114 (*sehA* ^Thr 95 Ala^, *sehB*), EN1115 (*sehA* ^Tyr 97 Ala^, *sehB*), and EN1116 (*sehA* ^Tyr 98 Ala^, *sehB*) strains were generated by removing Cm^R^ cassettes from EN1099, EN1100, EN1101, EN1102, EN1103, EN1104, EN1105, and EN1106, using plasmid pCP20 as described (Datsenko and Wanner, 2000).

### Ribosome profile analysis

Cells were grown for 2.5 h in N-minimal medium containing 0.01 mM Mg^2+^ and 10 mM arabinose and collected as described with modifications (Mohammad et al., 2019; Woolstenhulme et al., 2015). Cells were harvested by filtration using a Kontes 90 mm filtration apparatus with 0.45 μm nitrocellulose filters (Millipore, HAWP09000). During the filtration, cells on filter were scraped with a plastic spatula (SPL, 90040) and flash-frozen in liquid nitrogen. The pellets were grinded in the presence of 0.65 mL of lysis buffer (20 mM Tris pH 8.0, 10 mM MgCl_2_, 100 mM NH_4_Cl, 5 mM CaCl_2_, 0.1% NP-40, 0.4% Triton X-100, 100 U/mL DNase I, and 1 M NaCl) using SPEX 6775 freezer mill with 5 cycles of 1 min grinding at 5 Hz and 1 min of cooling. The lysates were cleared by centrifugation and then loaded onto 10 mL of 10-50% sucrose gradient buffer (20 mM Tris pH 8.0, 10 mM MgCl_2_, 100 mM NH_4_Cl, and 2 mM dithiothreitol (DTT)) and centrifuged in an SW-41 Ti rotor (Beckman) for 2 h at 40,000 rpm. Gradients were analyzed using an ISCO tube piercer (Brandel) and a liquid chromatography system equipped with an absorbance monitor (254 nm) and fraction collector (Bio-Rad).

### In vitro RNase III assays

For *in vitro* synthesis of shorter 16 rRNA transcripts, complementary oligonucleotides KU626 and KU627 containing T7 promoter were annealed. The annealed oligonucleotide was cloned into pBAD33 plasmid digested with SmaI. The cloned *rrsH* sequence retains 73 nt of the UTR sequence present at the 5’ and 3’ ends but is deleted for most of the mature 16S rRNA sequence. The pBAD33 plasmid harboring the stem structure of *rrsH* (corresponding to -131∼-102 and +1559∼+1589 relative to +1 of *rrsH* gene) under T7 promoter was linearized by digestion with XbaI (NEB, R0145).

Transcription of the linearized template is expected to contain the RNase III processing stem structure (30 and 31 nt at each side) and extra sequences derived from the plasmid (Δ16S-88nt). The T7 promoter-containing DNA template was transcribed using RiboMAX^TM^ Large Scale RNA Production Systems (Promega, P1300) according to the manufacturer’s instructions. Before reaction, each transcript in RNase III reaction buffer (10 mM Tris pH 7.9, 50 mM NaCl, 10 mM MgCl_2_, and 1 mM DTT; Invitrogen AM2290) was denatured at 95°C for 30 sec and immediately placed on ice. For RNase III digestion, ∼30 pmoles of *Salmonella* His_6_-tagged RNase III, His_6_-GFP-SehA, or His_6_-GFP-SehA^Y53A^ proteins were added in a final volume of 10 µl as indicated and incubated for 10 min at 37°C. Reactions were stopped by adding an equal volume of 2X TBU (7M urea, 12% Ficoll, and 1×TBE). The reaction products were analyzed by 15% denaturing TBE acrylamide gel containing 7 M urea and electrophoresed at 15 W for 25 min. Gels were stained using SYBR Gold (ThermoFisher, S11494) and visualized by UV detector (EMBI TEC, PI-1000).

### Northern blot analysis

To detect rRNA cleavage by SehA toxin *in vivo*, *Salmonella* strains harboring either the SehA-expressing plasmid or the empty vector were grown in N-minimal media containing 0.01 mM Mg^2+^. At OD_600_=0.1, cells were split into two flasks, one of which was induced by adding 10 mM L-arabinose and grown until vector-expressing cells reached OD_600_=0.3. Strains with the chromosomal mutations were grown similarly in N-minimal media containing 0.01 mM Mg^2+^ until wild-type *Salmonella* reached OD_600_=0.3 and extracted total RNA using Tri reagent (Ambion, 15596018). 10 µg of RNA samples were mixed to an equal volume of RNA sample loading buffer (Sigma Aldrich, R1386), incubated at 65°C for 5 min, and electrophoresed on a formaldehyde-agarose gel in 1 × MOPS buffer at 80 V for 30 min. The gel was transferred to nylon membrane (GE healthcare, Hybond-N hybridization membranes, GERPN303N) by capillary transfer for overnight. The transferred membrane was incubated with hybridization buffer (Invitrogen^TM^, ULTRAhyb^TM^-oligo, AM8663) at 42°C for 1 h, added 10^5^ cpm of labeled oligonucleotides specific for leader RNA or 16S ribosomal RNA sequences, and incubated for overnight. The membrane was washed at 42°C by low stringency washing buffer (Invitrogen^TM^, NorthernMax^TM^ Low Stringency Wash Buffer, AM8673) for 5 min twice and then washed at 42°C by high stringency washing buffer (Invitrogen^TM^, NorthernMax^TM^ High Stringency Wash Buffer, AM8674) for 15 min twice. The signal was detected using phosphorimager (Cytiva, Amersham^TM^ Typhoon Biomolecular Imager, 29187194).

### Electrophoretic mobility shift assay

300 ng of in vitro synthesized Δ16S-88 nt transcript was mixed with 0 to 5000 nM of purified GFP-tagged SehA proteins in binding buffer (50 mM Tris pH 7.5, 10 mM MgCl_2_, 250 mM KCl). The reactions were incubated at room temperature for 30 min in a total volume of 10 µL and stopped the reactions by adding loading dye (50% glycerol, 0.025% [wt/vol] bromophenol blue). Then, the samples were immediately resolved on 10% native polyacrylamide gel at 4°C in 0.5 × TBE buffer (45 mM Tris-HCl, 45 mM boric acid, 0.1 mM EDTA) for 1 h at 110 V. The gel was stained with SYBR Gold (Invitrogen, S33102) for 5 min and visualized by PrepOne^TM^ Sapphire Blue LED Gel Illuminator (EMBI TEC, PI-1000).

### Macrophage survival assay

Intramacrophage survival assays were conducted using the macrophage-like cell line J774 A.1. Briefly, 5 × 10^5^ macrophages in Dulbecco’s modified Eagle’s medium (DMEM) supplemented with 10% fetal bovine serum (FBS) were seeded in 24-well plates and cultured at 37°C. Overnight grown bacteria were added to the macrophages at a multiplicity of infection (MOI) of 10. The plates were centrifuged at 1000 rpm for 5 min at room temperature and incubated for an additional 20 min. Then, the extracellular bacteria were washed three times with PBS (phosphate-buffered saline) and killed by incubation with DMEM supplemented with 10% FBS and 120 µg mL^-1^ gentamicin for 1 h. For measuring the number of bacteria at 1 h, cells were lysed with PBS containing 0.1% Triton X-100 and plated on Luria-Bertani (LB) plates with appropriate dilutions. For measuring the number of bacteria at 21 h, the DMEM was replaced after 1 h with fresh DMEM containing 12 µg ml^-1^ gentamicin and the incubation was continued at 37°C. After 21 h, cells were lysed with PBS containing 0.1% Triton X-100 and plated on LB plates. The percentage survival was obtained by dividing the number of bacteria recovered after 21 h by the number of bacteria recovered at 1 h. All experiments were performed in triplicate and the results are representative of at least three independent experiments.

### Mouse virulence assay

Six-to eight-week-old female C3H/HeN mice were inoculated intraperitoneally with ∼10^3^ colony forming units of *Salmonella* strains. Mouse survival was followed for 21 days. Virulence assays were conducted three times with similar outcomes and the data correspond to groups of five mice. All animals were housed in a temperature- and humidity-controlled room, in which a 12 h light/12 h dark cycle was maintained. All procedures were performed according to the protocols (KW-181010-1) approved by the Institutional Animal Care and Use Committee of the Kangwon National University.

### Measuring non-replicating *Salmonella* inside macrophages

Macrophage infection and assessment of non-replicating *Salmonella* within macrophages were performed as described previously (Choi et al., 2019) with some modifications. *Salmonella* harboring mCherry-constitutive and GFP-inducible pFCcGi plasmid were grown overnight in Luria-Bertani medium with 10 mM L-arabinose to induce GFP expression and infect macrophages at a multiplicity of infection of 10. At 2 and 21 h postinfection, infected macrophages were washed and lysed with PBS solution containing 0.1 % Triton X-100 (Sigma) to release the intracellular bacteria. The fluorescence of the bacterial population was subsequently assessed by FACS analysis. Samples were analyzed on a NovoCyteTM Flow Cytometer (ACEA) using NovoExpress® software (ACEA). On the NovoCyteTM Flow Cytometer, fluorophores were excited at a wavelength of 488 nm, and green and red fluorescence were detected at 530 and 615 nm, respectively. Data were analyzed with NovoExpress® software. To analyze the fluorescence dilution, bacteria were identified after gating on the constitutive mCherry-positive signal. The percentage of cells expressing high levels of GFP (GFPHigh) inside J774 A.1 macrophages at the indicated times was calculated based on the following formula: The percentage of cells expressing high levels of GFP inside macrophages at the indicated times was calculated based on the following formula: (*Salmonella* above the grid)/(total number of *Salmonella*).

## DATA AVAILABILITY

All other relevant data are available from the corresponding author upon reasonable request.

## ACKNOWLEDGMENTS

This work was supported by Basic Science Research Program through the National Research Foundation of Korea (NRF) funded by the Ministry of Science, ICT and Future Planning [NRF-2022R1A2B5B02002256 and NRF-2020M3A9H5104235 to E.-J.L. and NRF-2021R1I1A1A01043879 to E.C.], the Ministry of Education (NRF-2020R1A6A3A13076438 to S.C.], and a grant from Korea University.

E.-J.L. designed the research, analyzed the data, and wrote the manuscript; S.C., Y.-J.C., and S.B. performed the experiments and wrote the manuscript; E.C. generated chromosomal mutant strains; Y.K.K. provided help with ribosome profiling experiment.

