## Supporting information for "A HigB-like toxin promotes non-replicating *Salmonella* inside macrophages by inhibiting ribonuclease III"

### SUPPLEMENTAL FIGURES

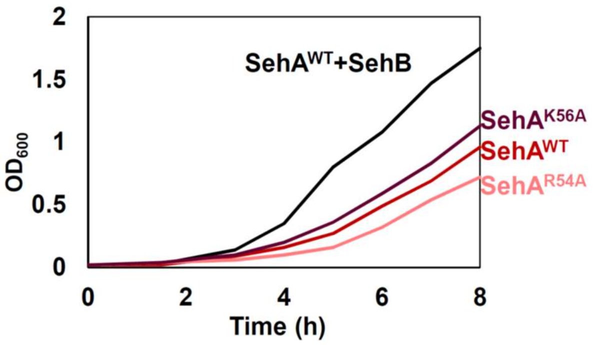


**Figure S1. Arg54 and Lys56 residues are not required for SehA toxin**
Growth curves of wild-type, the *sehB* deletion mutant (*sehB*Δ(155-429)) with the wild-type *sehA* gene or *sehA* derivatives with nucleotide substitutions (R54A or K56A) grown in N-minimal medium containing 10 mM $Mg^{2+}$ at 37°C for 8 h with shaking and measured absorbance at $OD_{600}$ every hour.

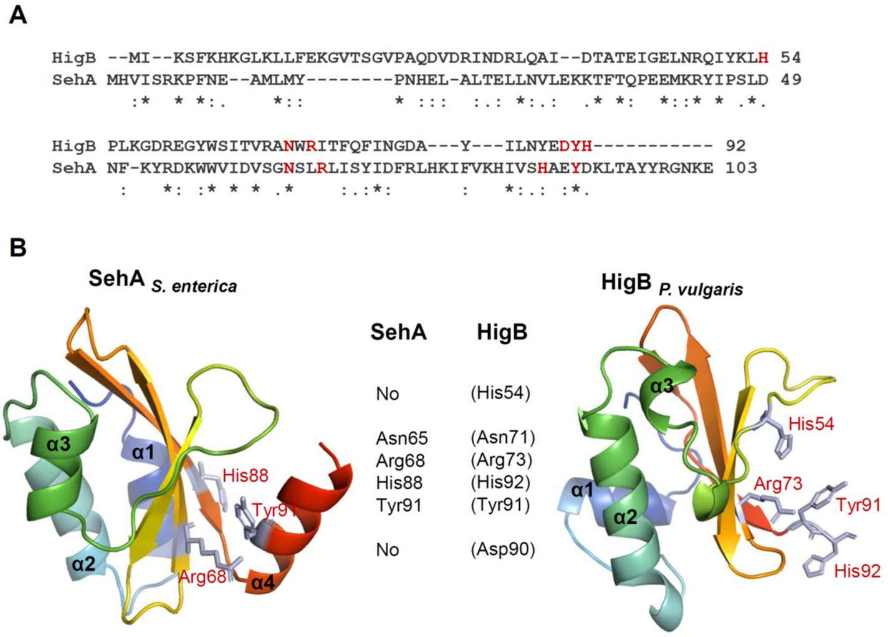


**Figure S2. SehA toxin contains an additional α4 helix compared to HigB *P. vulgaris***

(A) Sequence alignments of HigB from *Proteus vulgaris* and SehA toxin. The proposed active site residues in HigB and corresponding residues in SehA are colored in red. (B) Comparisons of active site residues (shown as sticks) in SehA and HigB *P. vulgaris* (PDB code 4MCT).

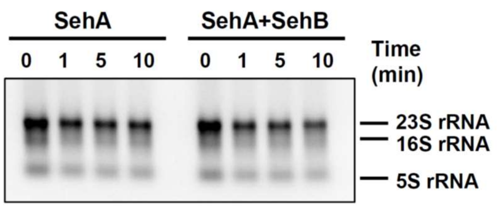


**Figure S3. SehA toxin alone does not exhibit endoribonuclease activity *in vitro***

Total RNAs were incubated with purified SehA proteins (left) or SehA preincubated with SehB proteins (right) for indicated times in room temperature. The reaction products were analyzed by MOPS-RNA gel.

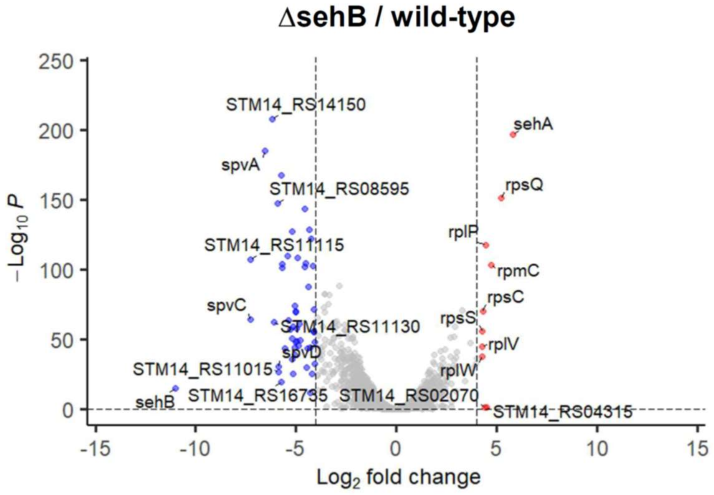


**Figure S4. SehA overexpression by deleting antitoxin SehB increases mRNA levels of ribosomal protein genes**

Volcano plot comparing  RNA-seq profiles of wild-type and *sehB* deletion mutant cells grown for 4 h in N-minimal medium containing 0.01 mM $Mg^{2+}$. Ten genes upregulated over 16-fold ($log_2$ fold > 4, right vertical dashed line) with adjusted p-value < 0.05 (horizontal dashed line) were labeled and dotted in red. 51 genes downregulated over 16-fold ($log_2$ fold < -4, left vertical dashed line) with adjusted p-value < 0.05 (horizontal dashed line) were dotted in blue. The 10 most upregulated or downregulated genes were labeled.

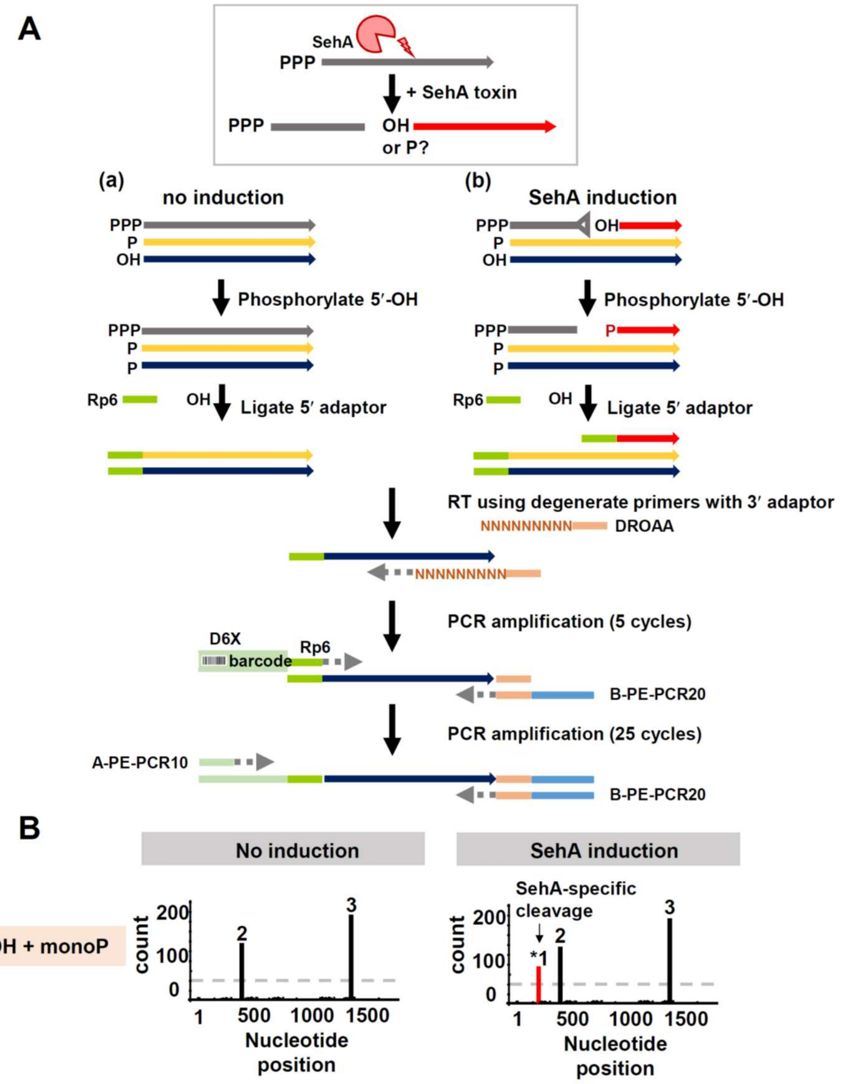


**Figure S5. Schematic representation of SehA-mediated RNA cleavage sequencing**

(A) Flowchart of SehA-mediated RNA cleavage sequencing. SehA-mediated endoribonucleolytic cleavage generates two RNA fragments, one of which might have either 5′-hydroxyl (5′-OH) or monophosphorylated (5′-monoP) groups. To generate cDNA libraries containing RNAs with 5′-OH or 5′-monoP groups, 5′-OH RNAs are phosphorylated by polynucleotide kinase (PNK). Then, the converted 5′-monophosphorylated RNAs are ligated with the Rp6 synthetic oligomer by T4 RNA ligase 1. After synthesizing cDNAs using degenerate primer with 3′ adaptor (DROAA), cDNAs from Rp6-ligated RNA are amplified by primers with the 5′-barcoded Rp6 (D6X) and 3′ adaptors (B-PE-PCR20). (B) Illumina sequencing identifies the first nucleotide of the Rp6-ligated cDNAs. Newly appeared reads upon SehA toxin induction (*1, shown in red line) correspond to the cleaved 5′-RNAs by SehA.

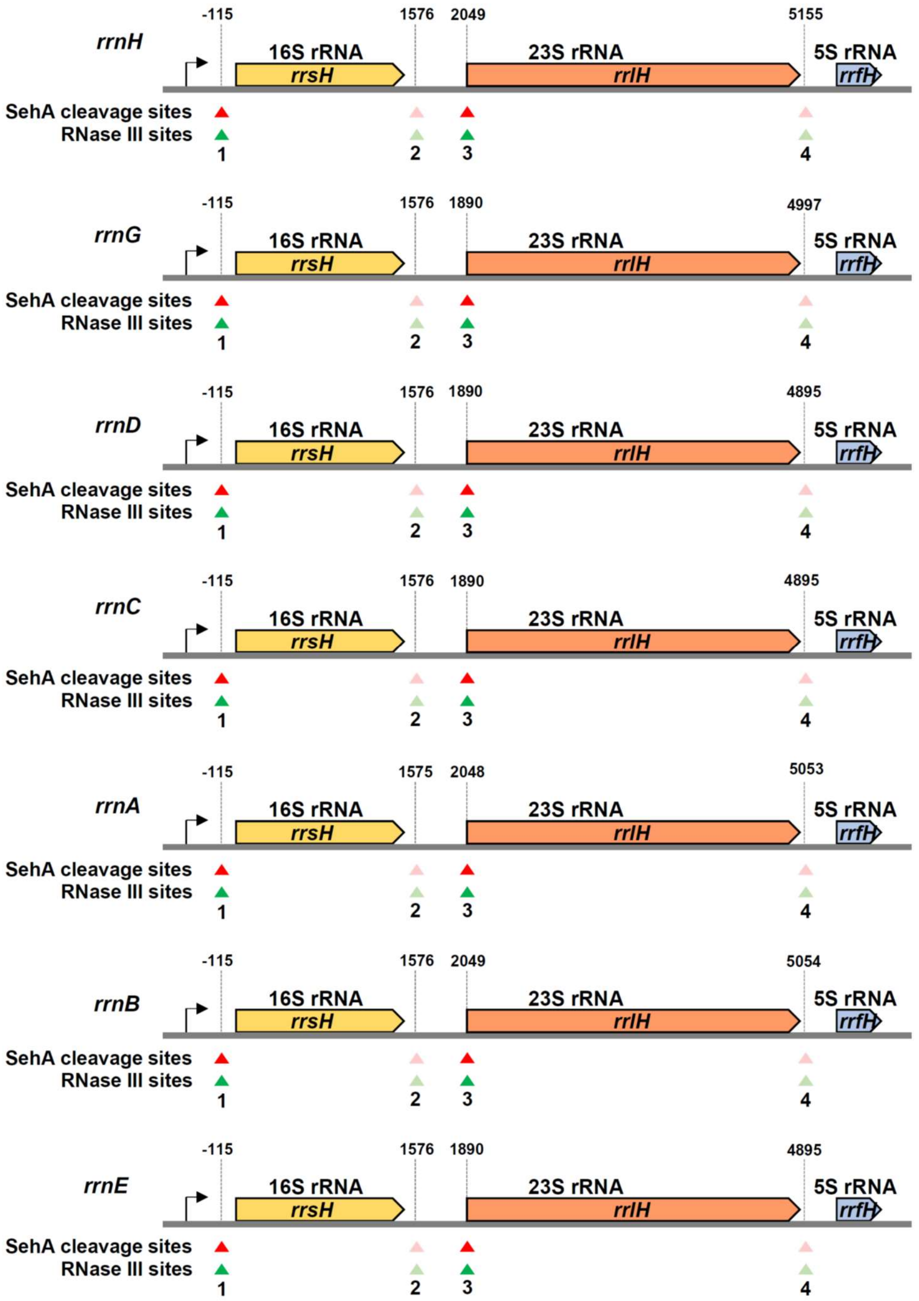


**Figure S6. SehA toxin cleavage sites are conserved in seven rRNA operons**

Schematic view of *Salmonella* 16S-23S-5S rRNA operon. Mature rRNA sequences are represented as colored boxes. The nucleotide positions are listed above relative to the 5' end of the mature 16S rRNA. SehA toxin cleavage sites are shown in red triangles and RNase III cleavage sites are shown in green triangles.

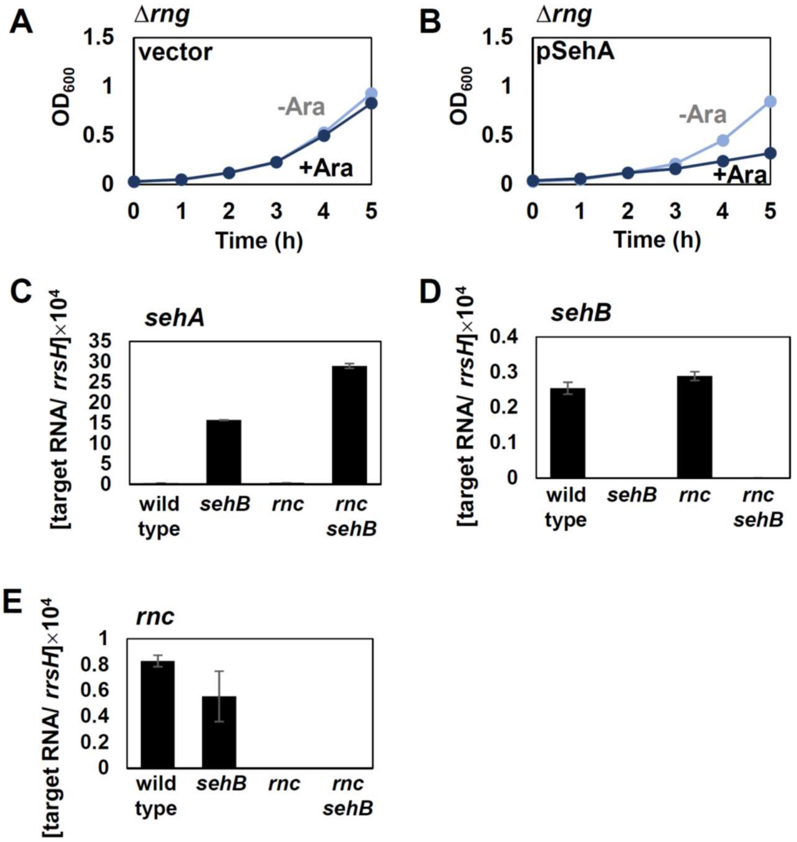


**Figure S7. SehA toxin is still effective in the *rng* mutant and *rnc* deletion does not affect *sehA* mRNA levels, related to Figure 6**

(A-B) Growth curves of the *rnc* deletion *Salmonella* strain expressing the empty vector (A) or pSehA (B) in N-minimal medium containing 10 mM $Mg^{2+}$ in the presence or absence of 1 mM L-arabinose. Bacteria were grown at 37°C for 5 h with shaking and measured absorbance at 600 nm every hour. (C-E) Relative mRNA levels of the *sehA* (A), *sehB* (B), and *rnc* (C) genes in *Salmonella* strains listed in Figure 6E.

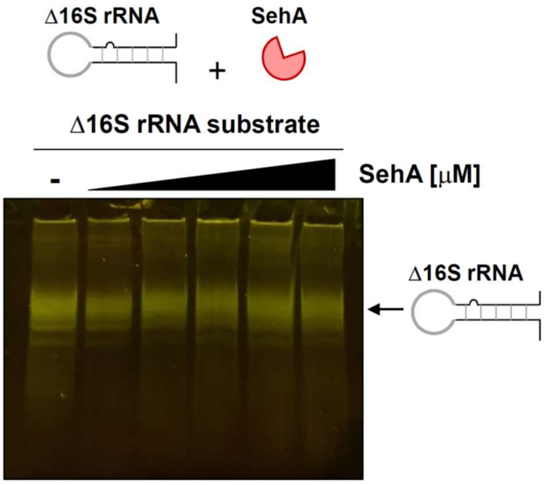


**Figure S8. SehA toxin does not bind to 16S rRNA *in vitro***

Gel mobility shift assay of 88 nt-long in vitro-synthesized 16S rRNA and purified GFP-tagged SehA toxin. 700 ng of in vitro-synthesized 16S rRNA was incubated with increasing concentrations of SehA toxin and electrophoresed as described in the Materials and Methods. Arrows mark the positions of the Δ16S rRNA. Concentrations were 0, 1000, 2000, 3000, 4000, and 5000 nM for lanes 1-6, respectively.

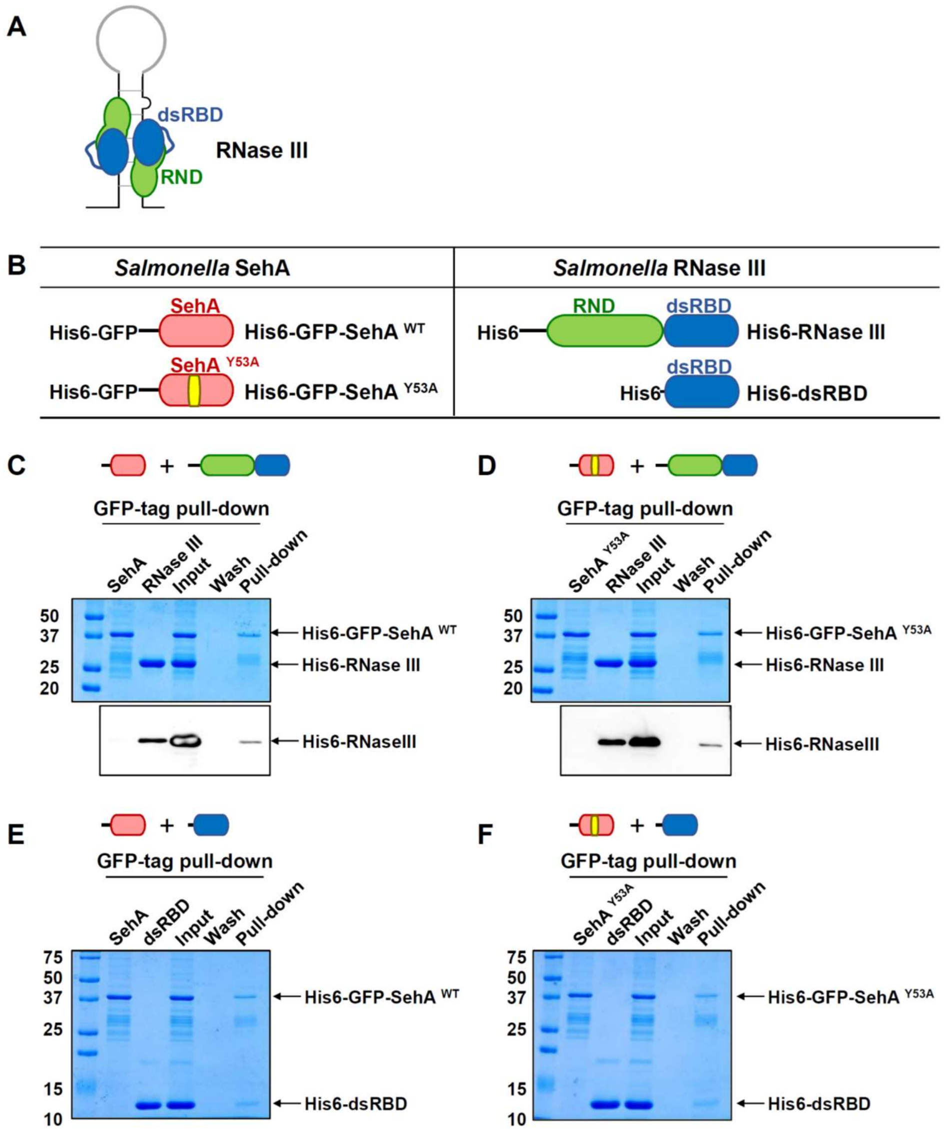


**Figure S9. SehA toxin binds to the dsRNA-binding domain of RNase III independently of toxin activity**

(A-B) Schematic representation of RNase III dimer-dsRNA complex (A) and RNase III and SehA proteins used in this experiment (B). (C-F) The GFP-tagged SehA or SehA Y53A bind to RNase III or the dsRNA-binding domain (dsRBD). (C-D) Pulldown assays of immobilized SehA WT (C) or SehA Y53A (D) in the presence of the full-length RNase III. The bands were detected with Coomassie blue staining or anti-His antibody. (E-F) Pulldown assays of immobilized SehA WT (E) or SehA Y53A (F) in the presence of the dsRNA-binding domain (dsRBD) of RNase III. The bands were detected with Coomassie blue staining.

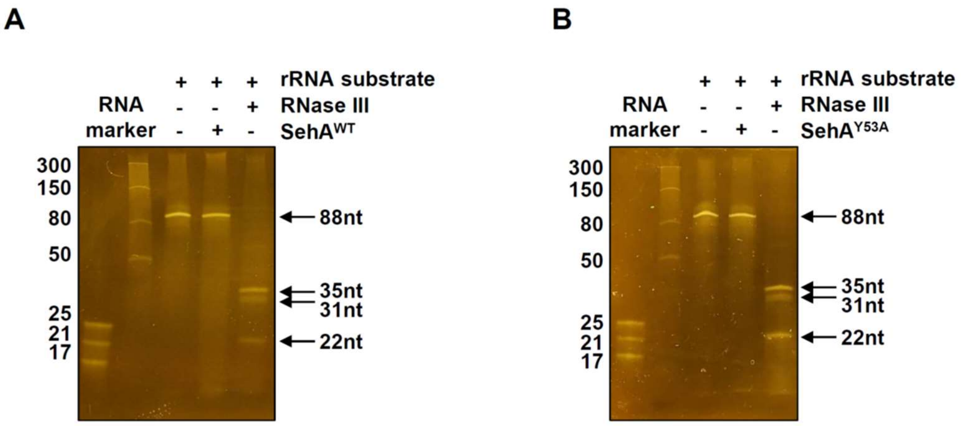


**Figure S10. *Salmonella* RNase III cleaves rRNA substrates, generating cleavage fragments *in vitro***

(A) Schematic diagram of *in vitro* synthesized 16S rRNA substrate. rRNA transcripts were synthesized in vitro, 88 nt RNA corresponding to sequence lacking most of the 16S rRNA mature sequence but retaining the stem structure flanking the 16S rRNA. (B) Sequence of DNA template for in vitro synthesized rRNA substrate. (C-D) In vitro RNase III cleavage assay of in vitro synthesized 16S rRNA substrate. The rRNA substrates were incubated with the *Salmonella* RNase III, $SehA^{WT}$, or $SehA^{Y53A}$. The reaction products were separated on a 15% TBE-urea gel and visualized by SYBR gold staining.

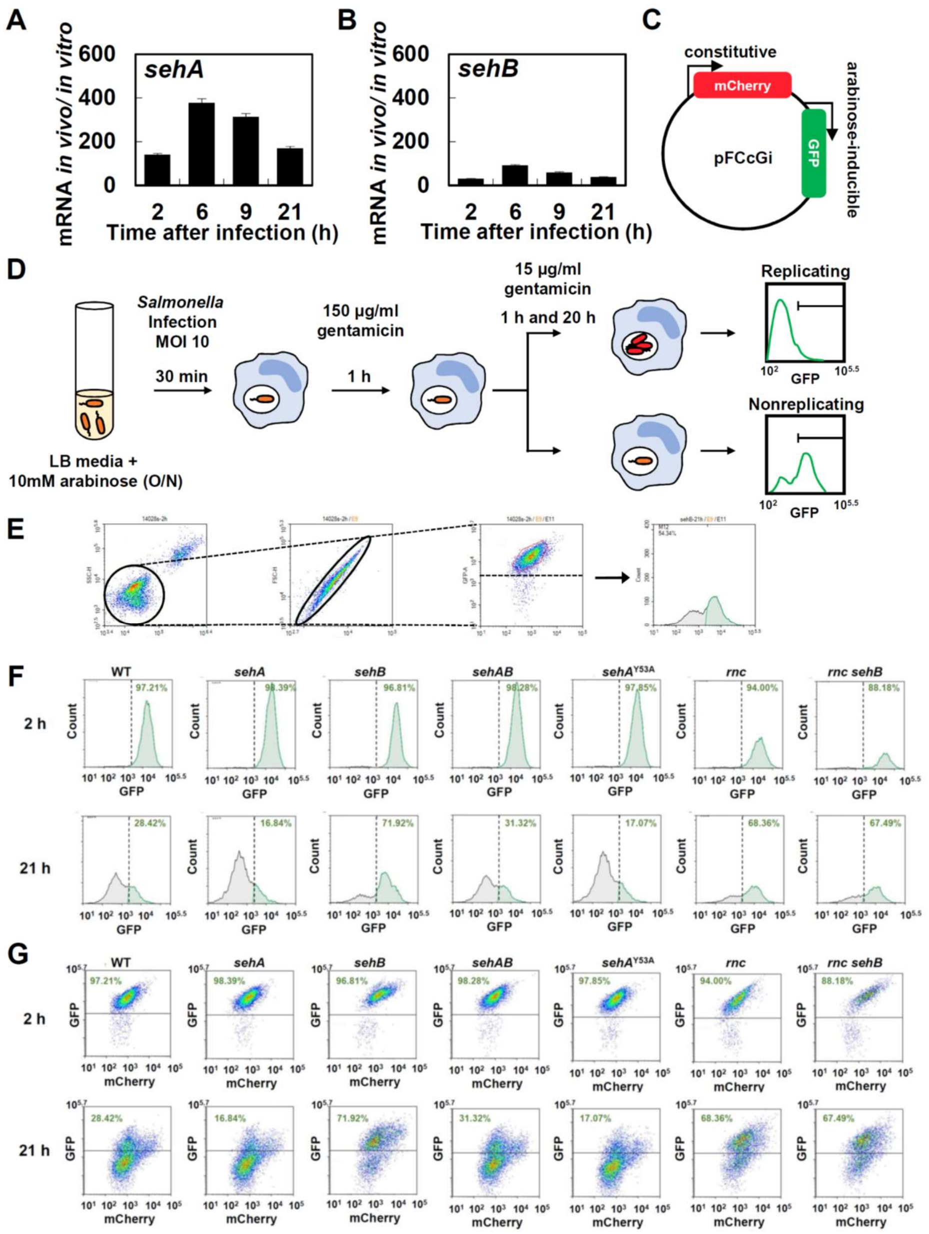


**Figure S11. SehA increases the formation of non-replicating *Salmonella* inside macrophages, related to Figure 8**

(A-B) Relative mRNA levels of the *sehA* (A) and *sehB* (B) genes produced by wild-type *Salmonella* (14028s) inside J774 A.1 macrophages at the indicated times after infection (n = 3 independent infections). *In vitro* mRNA levels were obtained from *Salmonella* grown in LB medium at early exponential growth phase and used for normalization. (C) Schematic representation of the pFCcGi plasmid. (D) Schematic representation of measuring non-replicating *Salmonella* inside macrophages by flow cytometry. (E) Gating strategy for flow cytometry analysis. Cells were firstly gated with FSC/SSC dot blot to select live bacteria, followed by second gating (FSC-H versus FSC-A) to select single cells. Then, GFP plots were analyzed after selecting bacteria that emit high levels of red fluorescence as shown in the right panel. (F) Histograms of *Salmonella* strains expressing GFP inside J774 A.1 macrophages. Histograms indicate the GFP fluorescence intensities of strains listed above at 2 or 21 h postinfection, with gates showing the fraction of the population exhibiting high levels of GFP. Non-replicating $GFP^{High}$ cells are indicated as green histograms. Bacterial fractions in the grid are indicated in green respectively. The percentage of cells expressing high levels of GFP inside macrophages at the indicated times was calculated based on the following formula: (*Salmonella* above the grid)/(total number of *Salmonella*). (G) Detection of mCherry and GFP fluorescence in *Salmonella* population harboring pFCcGi plasmid (n = 12,000 cells) inside J774 A.1 macrophages at 2 and 21 h postinfection in the absence of arabinose. Data are representative of four independent experiments.

Table S1. List of SehA-dependent cleavage site candidates in rRNA operons

| | | | | | | | OH+monoP | | monoP | | | OH+monoP | | monoP | |
|---|---|---|---|---|---|---|---|---|---|---|---|---|---|---|---|
| 열1 | Gene | Locus_tag | Product | Location | gbkey | Site 1 | value(-ara) | value(+ara) | value(-ara)3 | value(+ara)4 | Site 2 | value(-ara)2 | value(+ara)2 | value(-ara)4 | value(+ara)4 2 |
| *rrnH* | *rrsH* | STM14_0293 | 16s ribosomal RNA | 289879..291432 | rRNA | 289764 | 276.45 | 2116.24 | 1829 | 16505.81 | 291454 | 43.19 | 897.8 | 149 | 255.75 |
| | *rrlH* | STM14_0297 | 23S ribosomal RNA | 291925..295036 | rRNA | 291927 | 863.9 | 9170.36 | 12588 | 137449.52 | 295033 | 17.28 | 128.26 | 171 | 241.78 |
| | *rrfH* | STM14_0298 | 5S ribosomal RNA | 295221..295336 | rRNA | | | | | | | | | | |
| *rrnG* | *rrfG* | STM14_3254 | 5S ribosomal RNA | 2850278..2850393 (complement) | rRNA | | | | | | | | | | |
| | *rrlG* | STM14_3256 | 23S ribosomal RNA | 2850578..2853690 (complement) | rRNA | 2853688 | 1523.1 | 14762.67 | 30140.15 | 332403.12 | 2850581 | 50.77 | 492.09 | 940.38 | 996.15 |
| | *rrsG* | STM14_3258 | 16S ribosomal RNA | 2854026..2855577 (complement) | rRNA | 2855692 | 186.16 | 1722.31 | 3024.51 | 18318.82 | 2854002 | 186.16 | 1968.36 | 575.16 | 624.67 |
| *rrnF* | *rrfF* | STM14_4096 | 5S ribosomal RNA | 3580526..3580641 (complement) | rRNA | | | | | | | | | | |
| *rrnD* | *rrfD* | STM14_4098 | 5S ribosomal RNA | 3580771..3580886 (complement) | rRNA | | | | | | | | | | |
| | *rrlD* | STM14_4099 | 23S ribosomal RNA | 3580968..3583978 (complement) | rRNA | 3583976 | 1370.79 | 14885.7 | 30888.07 | 339414.06 | 3580971 | 135.36 | 615.11 | 1253.11 | 979.59 |
| | *rrsD* | STM14_4100 | 16S ribosomal RNA | 3584314..3585865 (complement) | rRNA | 3585980 | 795.4 | 6766.3 | 5572.28 | 43307.86 | 3584290 | 169.23 | 1845.33 | 577.35 | 605.74 |
| *rrnC* | *rrsC* | STM14_4686 | 16S ribosomal RNA | 4113825..4115376 | rRNA | 4113710 | 457.87 | 3719.45 | 2851 | 20313.1 | 4115400 | 69.11 | 1026.05 | 256 | 315.93 |
| | *rrlC* | STM14_4688 | 23S ribosomal RNA | 4115712..4118722 | rRNA | 4115714 | 760.23 | 8849.72 | 12616 | 139694.34 | 4118719 | 95.03 | 128.26 | 493 | 420.17 |

| | | | | | | | | | | | | | | | |
|---|---|---|---|---|---|---|---|---|---|---|---|---|---|---|---|
| | *rrfC* | STM14_4690 | 5S ribosomal RNA | 4118804..4118919 | rRNA | | | | | | | | | | |
| *rrnA* | *rrsA* | STM14_4794 | 16S ribosomal RNA | 4209750..4211301 | rRNA | 4209635 | 457.87 | 4553.12 | 2304 | 21229.73 | 4211325 | 257.17 | 1859.72 | 249 | 274.02 |
| | *rrlA* | STM14_4798 | 23S ribosomal RNA | 4211796..4214806 | rRNA | 4211798 | 881.18 | 10068.16 | 13088 | 143285.66 | 4214803 | 8.64 | 192.39 | 243 | 445.96 |
| | *rrfA* | STM14_4799 | 5S ribosomal RNA | 4214888..4215003 | rRNA | | | | | | | | | | |
| *rrnB* | *rrsB* | STM14_4967 | 16S ribosomal RNA | 4364715..4366266 | rRNA | 4364600 | 233.25 | 3334.68 | 2438 | 20375.43 | 4366290 | 95.03 | 1346.7 | 260 | 283.69 |
| | *rrlB* | STM14_4971 | 23S ribosomal RNA | 4366760..4369771 | rRNA | 4366763 | 786.15 | 8336.69 | 13091 | 143364.1 | 4369768 | 25.92 | 320.64 | 467 | 694.19 |
| | *rrfB* | STM14_4972 | 5S ribosomal RNA | 4369956..4370071 | rRNA | | | | | | | | | | |
| *rrnE* | *rrsE* | STM14_5021 | 16S ribosomal RNA | 4408256..4409807 | rRNA | 4408141 | 354.2 | 4617.24 | 2260 | 21398.35 | 4409831 | 25.92 | 256.51 | 241 | 296.59 |
| | *rrlE* | STM14_5023 | 23S ribosomal RNA | 4410143..4413153 | rRNA | 4410145 | 587.45 | 7374.76 | 12260 | 137263.62 | 4413150 | 25.92 | 256.51 | 340 | 381.48 |
| | *rrfE* | STM14_5024 | 5S ribosomal RNA | 4413338..4413378 | rRNA | | | | | | | | | | |

Table S2. List of SehA-dependent cleavage site candidates other than rRNA operons

| | | | | | | OH+monoP | | monoP | | | OH+monoP | | monoP | |
|---|---|---|---|---|---|---|---|---|---|---|---|---|---|---|
| **Gene** | **Locus_tag** | **Product** | **Location** | **gbkey** | **Site 1** | **value(-ara)** | **value(+ara)** | **value(-ara)3** | **value(+ara)3** | **Site 2** | **value(-ara)2** | **value(+ara)2** | **value(-ara)32** | **value(+ara)32** |
| STM14_RS23860 | STM14_5471 | tRNA_leu | 4823580..4823666 (complement) | tRNA | 4823666 | 101.54 | 861.16 | 8948.88 | 12145.51 | | | | | |
| *vapB* | STM14_RS16220 | TA system antitoxin VapB | 3214693..3214920 (complement) | CDS | 3214876 | 33.85 | 861.16 | 0 | 2.37 | | | | | |
| *secY* | STM14_4123 | preprotein translocase subunit SecY | 3599885..3601216 (complement) | CDS | 3600291 | 473.85 | 1353.25 | 6.56 | 4.73 | | | | | |
| *secG* | STM14_3976 | preprotein translocase subunit SecG | 3475036..3475368 (complement) | CDS | 3475452 | 0 | 984.18 | 13.12 | 82.82 | | | | | |
| FMN | | riboswitch | 3376863..3377061 (complement) | regulatory | 3377057 | 102 | 369.07 | 19.68 | 11.83 | 3377040 | 152.31 | 369.07 | 0 | 0 |
| *cpvA* | STM14_2910 | colicin V production protein | 2527443..2527931 (complement) | CDS | 2527955 | 33.85 | 615.11 | 2.19 | 4.73 | | | | | |
| *rnpB* | STM14_RS24250 | RNase P RNA component class A | 3433056..3433430 (complement) | ncRNA | 3433430 | 998.48 | 9472.72 | 15.31 | 70.98 | | | | | |
| STM14_RS13435 | STM14_2998 | cystein synthase B | 2603203..2604115 (complement) | CDS | 2603660 | 33.85 | 738.13 | 4.37 | 0 | | | | | |
| STM14_RS26105 | | hypothetical protein | 2814254..2815858 (complement) | CDS | 2814403 | 0 | 615.11 | 2.19 | 14.2 | | | | | |
| STM14_RS14630 | STM14_3281 | signal recognition particle protein | 2871119..2872481 (complement) | CDS | 2872510 | 16.92 | 738.13 | 0 | 4.73 | | | | | |
| *fliS* | STM14_2381 | flagellar export chaperone FliS | 2062225..2062632 | CDS | 2062345 | 4197 | 14393.61 | 0 | 0 | | | | | |
| STM14_RS24965 | STM14_2427 | helix-turn-helix domain-containing protein | 2093906..2094127 (complement) | CDS | 2094219 | 0 | 615.11 | 343.35 | 92.28 | | | | | |
| TPP | | riboswitch | 4396053..4396216 (complement) | regulatory | 127511 | 33.85 | 1107.2 | 61.23 | 182.19 | | | | | |
| *aceE* | STM14_0183 | pyruvate dehydrogenase (acetyl-transferring), homo dimeric type | 176953..179616 | CDS | 178400 | 50.77 | 861.16 | 0 | 1.07 | | | | | |

| | | | | | | | | | |
|---|---|---|---|---|---|---|---|---|---|
| *mntS* | STM14_974 | manganase accumulation protein MntS | 903009..903137 (complement) | CDS | 903188 | 744.63 | 2706.49 | 861.65 | 761.9 |
| STM14_0569 | STM14_0569 | DNase polymerase III subunit gamma/tau | 541601..543529 | CDS | 542538 | 50.77 | 615.11 | 0 | 0 |
| STM14_RS24185 | STM14_RS24185 | RtT sRNA | 1182719..1182854 (complement) | ncRNA | 1182850 | 16.92 | 492.09 | 0 | 0 |
| STM14_RS09910 | STM14_2181 | SpoVR family protein | 191499..1916531 | CDS | 1915936 | 0 | 2829.51 | 0 | 0 |

**Table S1.** Bacterial strains and plasmids used in this study

| Strain or plasmid | Description | Reference or source |
|---|---|---|
| ***S. enterica* serovar Typhimurium** | | |
| 14028s | wild-type | (Fields *et al.*, 1986) |
| MS7953s | *PhoP7953*::Tn10 | (Fields *et al.*, 1986) |
| EN1011 | *sehA*::$Cm^R$ | This study |
| EN1019 | *sehA* | This study |
| EN1098 | *sehB100*::$Cm^R$ | This study |
| EN1108 | *sehB100* | This study |
| EN1013 | *sehAB*::$Cm^R$ | This study |
| EN1021 | *sehAB* | This study |
| EN1050 | *sehA*::$tetRA^R$ | This study |
| EN1123 | *sehA*::$tetRA^R$/pKD46 | This study |
| EN1072 | *sehA* $^{Tyr\ 53\ Ala}$ | This study |
| EN1076 | *sehA* $^{Asp\ 61\ Ala}$ | This study |
| EN1045 | *sehA* $^{His\ 88\ Ala}$ | This study |
| EN1047 | *sehA* $^{Tyr\ 91\ Ala}$ | This study |
| EN1078 | *sehA* $^{lys\ 93\ Ala}$ | This study |
| EN1081 | *sehA* $^{Thr\ 95\ Ala}$ | This study |
| EN1063 | *sehA* $^{Tyr\ 97\ Ala}$ | This study |
| EN1084 | *sehA* $^{Tyr\ 98\ Ala}$ | This study |
| EN1099 | *sehA* $^{Tyr\ 53\ Ala}$, *sehB100*::$Cm^R$ | This study |
| EN1100 | *sehA* $^{Asp\ 61\ Ala}$, *sehB100*::$Cm^R$ | This study |
| EN1101 | *sehA* $^{His\ 88\ Ala}$, *sehB100*::$Cm^R$ | This study |
| EN1102 | *sehA* $^{Tyr\ 91\ Ala}$, *sehB100*::$Cm^R$ | This study |
| EN1103 | *sehA* $^{lys\ 93\ Ala}$, *sehB100*::$Cm^R$ | This study |
| EN1104 | *sehA* $^{Thr\ 95\ Ala}$, *sehB100*::$Cm^R$ | This study |
| EN1105 | *sehA* $^{Tyr\ 97\ Ala}$, *sehB100*::$Cm^R$ | This study |
| EN1106 | *sehA* $^{Tyr\ 98\ Ala}$, *sehB100*::$Cm^R$ | This study |
| EN1109 | *sehA* $^{Tyr\ 53\ Ala}$, *sehB100* | This study |
| EN1110 | *sehA* $^{Asp\ 61\ Ala}$, *sehB100* | This study |
| EN1111 | *sehA* $^{His\ 88\ Ala}$, *sehB100* | This study |

| | | |
|---|---|---|
| EN1112 | *sehA* $^{Tyr\ 91\ Ala}$, *sehB100* | This study |
| EN1113 | *sehA* $^{lys\ 93\ Ala}$, *sehB100* | This study |
| EN1114 | *sehA* $^{Thr\ 95\ Ala}$, *sehB100* | This study |
| EN1115 | *sehA* $^{Tyr\ 97\ Ala}$, *sehB100* | This study |
| EN1116 | *sehA* $^{Tyr\ 98\ Ala}$, *sehB100* | This study |
| SM368 | *rnc*::Cm$^{R}$ | This study |
| SM371 | *rnc* | This study |
| SM376 | *rnc, sehB*::Cm$^{R}$ | This study |
| SM383 | *rnc, sehB* | This study |
| YT159 | 14028s/pBAD33 | This study |
| MJ003 | 14028s/pBAD33-*sehA* | This study |
| SM321 | 14028s/pBAD33-*sehA* $^{Tyr\ 53\ Ala}$ | This study |
| SM373 | *rnc*/pBAD33 | This study |
| SM378 | *rnc*/pBAD33-*sehA* | This study |
| SM374 | *rnc*/pBAD33-*rnc* | This study |
| SM299 | 14028s/pFCcGi | (Choi *et al*., 2019) |
| SW59 | *sehA*/pFCcGi | This study |
| SM154 | *sehB*/pFCcGi | This study |
| SW62 | *sehA* $^{Tyr\ 53\ Ala}$ /pFCcGi | This study |
| SM392 | *rnc*/pFCcGi | This study |
| SM393 | *rnc, sehB*/pFCcGi | This study |
| SM483 | DH5α/pBAD33-T7-*rrsH* leader | This study |
| ***Escherichia coli K-12*** | | |
| DH5α | *fhuA2 lac(del)U169 phoA glnV44 Φ80' lacZ(del)M15 gyrA96 recA1 relA1 endA1 thi-1 hsdR17* | (Taylor *et al,* 1993*)* |
| MJ001 | DH5α/pBAD33-*sehA* | This study |
| SM317 | DH5α/pBAD33-*sehA* $^{Tyr\ 53\ Ala}$ | This study |
| SM316 | DH5α/pBAD33-*rnc* | This study |
| SM386 | DH5α/pRSF-sfGFP-*sehA* | This study |
| SW112 | DH5α/ pRSF-sfGFP-*sehA* $^{Tyr\ 53\ Ala}$ | This study |
| JH001 | DH5α/pRSF-sfGFP-*sehB* | This study |
| SM461 | DH5α/Gst-YA-*sehA* | This study |
| SM397 | DH5α/Gst-YA-*sehA* $^{Tyr\ 53\ Ala}$ | This study |
| SM361 | DH5α/pET-His-FP-*rnc* | This study |

| | | |
|---|---|---|
| SM462 | DH5α/pET-His-FP-rnc dsRNA binding domain (155-227 aa) | This study |
| SM463 | DH5α/pET-His-FP-rnc RNase domain (1-128 aa) | This study |
| ***Escherichia coli K-12*** | | |
| BL21(DE3) | *B F–, ompT, gal, dcm, lon, hsdS$_B$(r$_B$– m$_B$–), λ(DE3 [lacI lacUV5-T7p07 ind1 sam7 nin5]), [malB$^+$]$_{K-12}$(λ$^S$)* | (Wood. W. B., 1966) |
| SM394 | BL21/pRSF-sfGFP-*sehA* | This study |
| SM395 | BL21/ pRSF-sfGFP-*sehA* $^{Tyr\ 53\ Ala}$ | This study |
| JH002 | BL21/pRSF-sfGFP-*sehB* | This study |
| SM365 | BL21/pET-His-FP-*rnc* | This study |
| SM464 | BL21/pET-His-FP-rnc dsRNA binding domain | This study |
| SM465 | BL21/pET-His-FP-rnc RNase domain | This study |
| **plasmids** | | |
| pBAD33 | pACYC184 *ori* Cm$^r$ | (Guzman *et al,* 1995) |
| pRSF-sfGFP | T7promoter TEV RSF*ori dam*$^-$ Km$^r$ | (Sung & Song, 2014) |
| pET-His-FP | T7promoter TEV ColEI*ori* Km$^r$ | (Kim *et al.*, 2013) |
| pKD3 | repR$_{6K\gamma}$ Ap$^R$ FRT Cm$^R$ FRT | (Datsenko & Wanner, 2000) |
| pKD46 | rep$_{pSC101}$$^{ts}$ Ap$^R$ P$_{araBAD}$ γ β exo | (Datsenko & Wanner, 2000) |
| pCP20 | rep$_{pSC101}$$^{ts}$ Ap$^R$ Cm$^R$ *cI857* λP$_R$*flp* | (Datsenko & Wanner, 2000) |
| pFCcGi | *rpsM*::*mCherry* and PBAD::*gfpmut3a* promoter fusions in pFPV25.1, Ap$^R$ | (Figueira R *et al,* 2013) |
| pBAD33-T7-*rrsH* leader | *rrsH* leader (Δ16S-88nt) | This study |


Datsenko, K.A., and Wanner, B.L. (2000) One-step inactivation of chromosomal genes in *Escherichia coli* K-12 using PCR products. *Proc Natl Acad Sci U S A* **97**: 6640-6645.

Fields, P.I., Swanson, R.V., Haidaris, C.G., and Heffron, F. (1986) Mutants of *Salmonella typhimurium* that cannot survive within the macrophage are avirulent. *Proc Natl Acad Sci U S A* **83**: 5189-5193.

Kim, B.W., Hong, S.B., Kim, J.H., Kwon, D.H., and Song, H.K. (2013) Structural basis for recognition of autophagic receptor NDP52 by the sugar receptor galectin-8. *Nat Commun* **4**: 1613.

Sung, K.H., and Song, H.K. (2014) Direct recognition of the C-terminal polylysine residues of nonstop protein by Ltn1, an E3 ubiquitin ligase. *Biochem Biophys Res Commun* **453**: 642-647.

**Table S2.** Primers used in this study.

| Name | Sequence (from 5' to 3') | |
|---|---|---|
| **Knockout or deletion** | | |
| KHU634 | AATTGTTTTCAGGTTGAAAACTTTGTAATGGAACTACATTTG TAGGCTGGAGCTGCTTCG | *sehA* deletion $Cm^R$ cassette insertion |
| KHU635 | TTTTTTGCGCTGGTTGCATCCATCTGACGATGAGTTCTCAC ATATGAATATCCTCCTTAG | *sehA* deletion $Cm^R$ cassette insertion |
| KHU637 | AAGAGGGCGGGAATGAGGAGGAGTAGCTGGACATATCCA ACATATGAATATCCTCCTTAG | *sehB::100* deletion $Cm^R$ cassette insertion |
| KHU944 | GTGGAGTTTCTGGTCGACCACGACGATCTTGAAAACCCAC TGTAGGCTGGAGCTGCTTCG | *sehB::100* deletion $Cm^R$ cassette insertion |
| KHU694 | AATTGTTTTCAGGTTGAAAACTTTGTAATGGAACTACATTTT AAGACCCACTTTCACATTTAAG | *sehA* $Tet^R$ cassette insertion |
| KHU695 | ATCTTTTTTGCGCTGGTTGCATCCATCTGACGATGAGTTCC TAAGCACTTGTCTCCTGTTTAC | *sehA* $Tet^R$ cassette insertion |
| KHU654 | TTGAAAACTGGATAACGTTGCGCTA | *sehA* mutagenesis product insertion |
| KHU700 | AACCAAGTTCGTTGGCAAGATCTGC | *sehA* mutagenesis product insertion |
| KU288 | CGGAAGCTGGGCTACACTTTTAATCATCAGGAGCTGTTGC TGTAGGCTGGAGCTGCTTCG | *rnc* deletion $Cm^R$ cassette insertion |
| KU289 | TCCTGATCGTGCGCCTCGCCGCGTACCTGCACAACCAGG TCATATGAATATCCTCCTTAG | *rnc* deletion $Cm^R$ cassette insertion |
| **Site-directed mutagenesis** | | |
| KHU752 | AATTTTAAGGCCAGAGATAAA | *sehA* $^{Y53A}$ chromosomal mutation |
| KHU753 | TTTATCTCTGGCCTTAAAATT | *sehA* $^{Y53A}$ chromosomal mutation |
| KHU754 | TGGGTGATTGCTGTTTCTGGT | *sehA* $^{D61A}$ chromosomal mutation |
| KHU755 | ACCAGAAACAGCAATCACCCA | *sehA* $^{D61A}$ chromosomal mutation |
| KHU690 | ATCGTTTCCGCTGCCGAATAT | *sehA* $^{M88A}$ chromosomal mutation |
| KHU691 | ATATTCGGCAGCGGAAACGAT | *sehA* $^{M88A}$ chromosomal mutation |
| KHU692 | CATGCCGAAGCTGACAAACTG | *sehA* $^{Y91A}$ chromosomal mutation |
| KHU693 | CAGTTTGTCAGCTTCGGCATG | *sehA* $^{Y91A}$ chromosomal mutation |
| KHU713 | GAATATGACGCACTGACCGCA | *sehA* $^{K93A}$ chromosomal mutation |
| KHU714 | TGCGGTCAGTGCGTCATATTC | *sehA* $^{K93A}$ chromosomal mutation |
| KHU715 | GACAAACTGGCCGCATACTAT | *sehA* $^{T95A}$ chromosomal mutation |
| KHU716 | ATAGTATGCGGCCAGTTTGTC | *sehA* $^{T95A}$ chromosomal mutation |
| KHU717 | CTGACCGCAGCCTATCGGGGT | *sehA* $^{Y97A}$ chromosomal mutation |
| KHU718 | ACCCCGATAGGCTGCGGTCAG | *sehA* $^{Y97A}$ chromosomal mutation |
| KHU719 | ACCGCATACGCTCGGGGTAAT | *sehA* $^{Y98A}$ chromosomal mutation |
| KHU720 | ATTACCCCGAGCGTATGCGGT | *sehA* $^{Y98A}$ chromosomal mutation |
| **Cloning** | | |
| KHU859 | GCTCTAGACTTTGTAATGGAACTACATTGTGCATGTTA TCAGCCGAAA | *sehA*-Xba1 |

| | | |
|---|---|---|
| KHU860 | CCCAAGCTTTCATTCTTTATTACCCCGAT | *sehA*-Hindiii |
| KU280 | GCTCTAGACATTCATTTATTGGTATCGCATGAACCCCA TCGTAATTAA | *rnc*-Xba1 |
| KU281 | CCCAAGCTTTCATTCCAACTCCAGTTTT | *rnc*-Hindiii |
| KU285 | CGGGATCCGAACCCCATCGTAATTAATCG | *rnc*-BamH1- pET |
| KHU897 | CGCGGATCCGTGCATGTTATCAGCCGA | *sehA*-BamH1 |
| KHU898 | CCGCTCGAGTCATTCTTTATTACCCCG | *sehA*-Xho1 |
| KHU899 | CGCGGATCCATGGATGCAACCAGCGCA | *sehB*-BamH1 |
| KHU900 | CCGCTCGAGCTACTCGATGAAGGCATC | *sehB*-Xho1 |
| KU287 | CCGCTCGAGTCATTCCAACTCCAGTTTTT | *rnc*-Xho1 |
| KU551 | CGCGGATCCCCGAAAACGCGCTTGCAGGA | *rnc dsRNA binding*-BamH1 |
| KU552 | CCGCTCGAGTCAATTACTATCGAGAAATAC | *Rnc RNase*-Xho1 |
| **OH-cleavage sequencing** | | |
| Rp6 | CGGCACCAACCGAGGVVVVVVVCGC | 5' ligation adapter |
| DROAA | GGCATTCCTGCTGAACCGCTCTTCCGATCTNNNNNNNNAA | Reverse transcription |
| D6A | CTCTTTCCCTACACGACGCTCTTCCGATCTNTACACGGCA CCAACCGAGG | Barcode A |
| D6B | CTCTTTCCCTACACGACGCTCTTCCGATCTNGTATCGGCA CCAACCGAGG | Barcode B |
| D6C | CTCTTTCCCTACACGACGCTCTTCCGATCTNCGTCCGGCA CCAACCGAGG | Barcode C |
| D6D | CTCTTTCCCTACACGACGCTCTTCCGATCTNAAGTCGGCA CCAACCGAGG | Barcode D |
| D6E | CTCTTTCCCTACACGACGCTCTTCCGATCTNACACCGGCA CCAACCGAGG | Barcode E |
| A-PE-PCR10 | AATGATACGGCGACCACCGAGATCTACACTCTTTCCCTAC ACGACG | Sequencing prep PCR |
| B-PE-PCR20 | CAAGCAGAAGACGGCATACGAGATCGGTCTCGGCATTCC TGCTGAACCGC | Sequencing prep PCR |
| **Northern blot analysis** | | |
| NB_probe-L1 | CAGTGCAACGCGGCTTTCGC | Probe for 16S rRNA leader |
| NB_probe-L2 | TTCGTCCGAGGACGTTAAGA | Probe for 16S rRNA leader |
| NB_probe-R3 | ATGGCTGCATCAGGCTTGCG | Probe for 16S rRNA |
| NB_probe-R4 | TATTAACCACAACACCTTCC | Probe for 16S rRNA |
| ***In vitro* RNase III assay** | | |
| KU626 | TAATACGACTCACTATAGGACAATTTATCAGACAATCTGTGT GGGCACTCGAAGGGGTTTGCAGTGCTCACACAGATTGTC TGATGAAAAAC | *rrsH* 5' UTR annealing-F |
| KU627 | GTTTTTCATCAGACAATCTGTGTGAGCACTGCAAACCCCTT CGAGTGCCCACACAGATTGTCTGATAAATTGTCCTATAGTG AGTCGTATTA | *rrsH* 5' UTR annealing-R |
| **qRT PCR** | | |
| KHQ045 | TTCAATGAAGCGATGCTCATG | *sehA* qRT PCR |
| KHQ046 | GAGTGAATGTTTTTTTCTCCAGAACA | *sehA* qRT PCR |
| KHQ047 | ATGCAACCAGCGCAAAAAAG | *sehB* qRT PCR |
| KHQ048 | GCCCTGCGATACTCATTGTCA | *sehB* qRT PCR |
| KHQ143 | AGTCCGGGCGATAAACAAAA | *rnc* qRT PCR |
| KHQ144 | CGCGTACCTGCACAACCA | *rnc* qRT PCR |
| 6970 | CCAGCAGCCGCGGTAAT | *rrsH* qRT PCR |
| 6971 | TTTACGCCCAGTAATTCCGATT | *rrsH* qRT PCR |
